# Ensemble-level organization of human kinetochores and evidence for distinct tension and attachment sensors

**DOI:** 10.1101/685248

**Authors:** Emanuele Roscioli, Tsvetelina E. Germanova, Christopher A. Smith, Peter A. Embacher, Muriel Erent, Amelia I. Thompson, Nigel J. Burroughs, Andrew D. McAinsh

## Abstract

Kinetochores are multi-protein machines that form dynamic attachments to microtubules and generate the forces for chromosome segregation. High-fidelity is ensured because kinetochores can monitor attachment status and tension, using this information to activate checkpoints and error correction mechanisms. To explore how kinetochores achieve this we used two and three colour subpixel fluorescence localisation to define how six protein subunits from the major kinetochore complexes CCAN, MIS12, NDC80, KNL1, RZZ and the checkpoint proteins Bub1 and Mad2 are organised in the human kinetochore. This reveals how the kinetochore outer plate is a liquid crystal-like system with high nematic order and largely invariant to loss of attachment or tension except for two mechanical sensors. Firstly, Knl1 unravelling relays tension and secondly NDC80 jack-knifes under microtubule detachment, with only the latter wired up to the checkpoint signalling system. This provides insight into how kinetochores integrate mechanical signals to promote error-free chromosome segregation.

## Introduction

Human kinetochores are multi-megadalton sized protein machines that assemble on the centromeres of every sister chromatid and provide an attachment site for the tips of ~20 dynamic spindle microtubules (that comprise the kinetochore (K)-fibre). Kinetochores must coordinate microtubule dynamics within the K-fibre and maintain attachment during phases of growth and shrinkage – thus coupling the energy release from microtubule depolymerisation to do work (Auckland and McAinsh, 2015; Rago and Cheeseman, 2013). These kinetochore-microtubule attachments are essential for the accurate segregation of chromosomes in all eukaryotes. There is however limited understanding of how this machinery adapts to changes in microtubule occupancy and/or the imposition of pushing and pulling forces. These are important questions because a substantial body of work indicates that kinetochores must be able to sense changes in tension and occupancy (Long et al., 2019), sensors that underpin decision making and error correction of the kinetochore.

Classic biophysical experiments established how applying tension to a kinetochore stabilised the attachment to microtubules (Nicklas and Koch, 1969) and that this was coupled to changes in the chemical (phosphorylation) state of the kinetochore (Nicklas et al., 1995). Live imaging experiments further show that tension between sister kinetochores (as measured by changes in inter-kinetochore (K-K) distance) can explain the oscillatory movements of bi-orientated kinetochores, being a determinant of directional switching (Burroughs et al., 2015; Wan et al., 2012). Changes in the K-K distance were also implicated in error correction processes that destabilise improper kinetochore attachments and stabilise bi-orientation (Lampson and Cheeseman, 2011). Recent work has however shown how low K-K tension kinetochores do not necessarily trigger error correction (Dudka et al., 2018). Furthermore, the imposition of K-K tension does not appear to be required for silencing of the spindle assembly checkpoint (SAC). By correlating the number of kinetochore-bound microtubules with checkpoint protein recruitment it looks like the crucial transition to SAC silencing occurs at around half-maximal occupancy (Dudka et al., 2018; Etemad et al., 2015; Etemad et al., 2019; Kuhn and Dumont, 2017). Kinetochores thus appear to be able to “count” the number of bound microtubules.

Intra-kinetochore tension may generate key mechanical signals that are sensitive to changes in microtubule attachment and/or the imposition of force (for review see: (Maresca and Salmon, 2010)). Intra-kinetochore tension refers to the measurement of changes in distance between two components of the kinetochore labelled with different fluorophores (denoted by delta, Δ). Initial pioneering experiments showed that increased Δ and not the K-K tension correlates with SAC silencing (Maresca and Salmon, 2009; Uchida et al., 2009; Wan et al., 2009). However, there is also evidence that high Δ is not always necessary for SAC silencing (Etemad et al., 2015; Magidson et al., 2016; Tauchman et al., 2015). It thus remains uncertain how the kinetochore monitors changes in occupancy and whether it can sense changes in tension at all. One idea is that there are kinetochore conformations or tensile elements that would function as tension and/or attachment sensors. While X-ray crystallography and electron microscopy are beginning to provide a structural view of the kinetochore (Hamilton et al., 2019; Pesenti et al., 2016; Welburn and Cheeseman, 2008) this approach is limited to subsets of kinetochore components and does not allow the impact of microtubule binding and forces to be easily determined. Furthermore, these approaches are limited to single assembles, while the human kinetochore in a living cell incorporates multiple microtubule attachment sites (~20) populated with multiple copies of each kinetochore complex (Huis In ‘t Veld et al., 2016; Johnston et al., 2010; Suzuki et al., 2011). Thus, the higher-order, ensemble-level organisation of the human kinetochore remains out of reach. Here, we developed experimental and computational methods to investigate changes in kinetochore architecture within an intact cell and integrated this information with distances and geometry defined in structural studies. We characterized the ensemble organization of the inner-outer kinetochore and found that Ndc80 and Knl1 are crucial elements that sense microtubule attachment and tension, respectively.

## Results

### Measurement of 3D Euclidian distances between kinetochore proteins

To obtain insight into the 3D nanoscale architecture of the human kinetochore we deployed an image acquisition and computational pipeline that outputs the 3D Euclidian distances (Δ_3D_) between differentially labelled kinetochore proteins in near-diploid hTERT-RPE1 cells (from now on referred as RPE1; (Smith et al., 2016). One limitation of this approach is the overestimation of mean distances (Suzuki et al., 2018). This is because Euclidian distances cannot be negative leading to a positive bias in the Δ_3D_ distribution, *i.e*. distances are overestimated (Figure 1A; Supplemental Table 1). To correct for this bias we developed an algorithm to infer the true Euclidian distances (Δ_EC_) between two fluorophores. This algorithm goes beyond previous methods (Churchman et al., 2005; Suzuki et al., 2018) by taking into account the non-anisotropic point spread function in 3D datasets (see Supplemental Methods for details). To test the accuracy of this method we fixed cells with 4% formaldehyde/0.2% triton and stained them with an anti-CenpC antibody that recognises the amino-terminal region of the protein (amino acids 1-426), binding to a site preceding the region binding CenpA nucleosomes. This primary antibody was then detected using a mixture of three different secondary antibodies (conjugated to Alexa Fluor 488, 568 and 647; Figure 1A). The Euclidean bias corrected delta distances were 2.2±1.5nm (mean ± standard deviation; n=2302; 488-to-568), 4.3±2.7nm (n=1437; 488-to-647) and 4.5±2.8nm (n=1452; 568-to-647), considerably smaller than the corresponding Δ_3D_ (Figure 1A; Supplemental Table 1).

**Figure 1.**
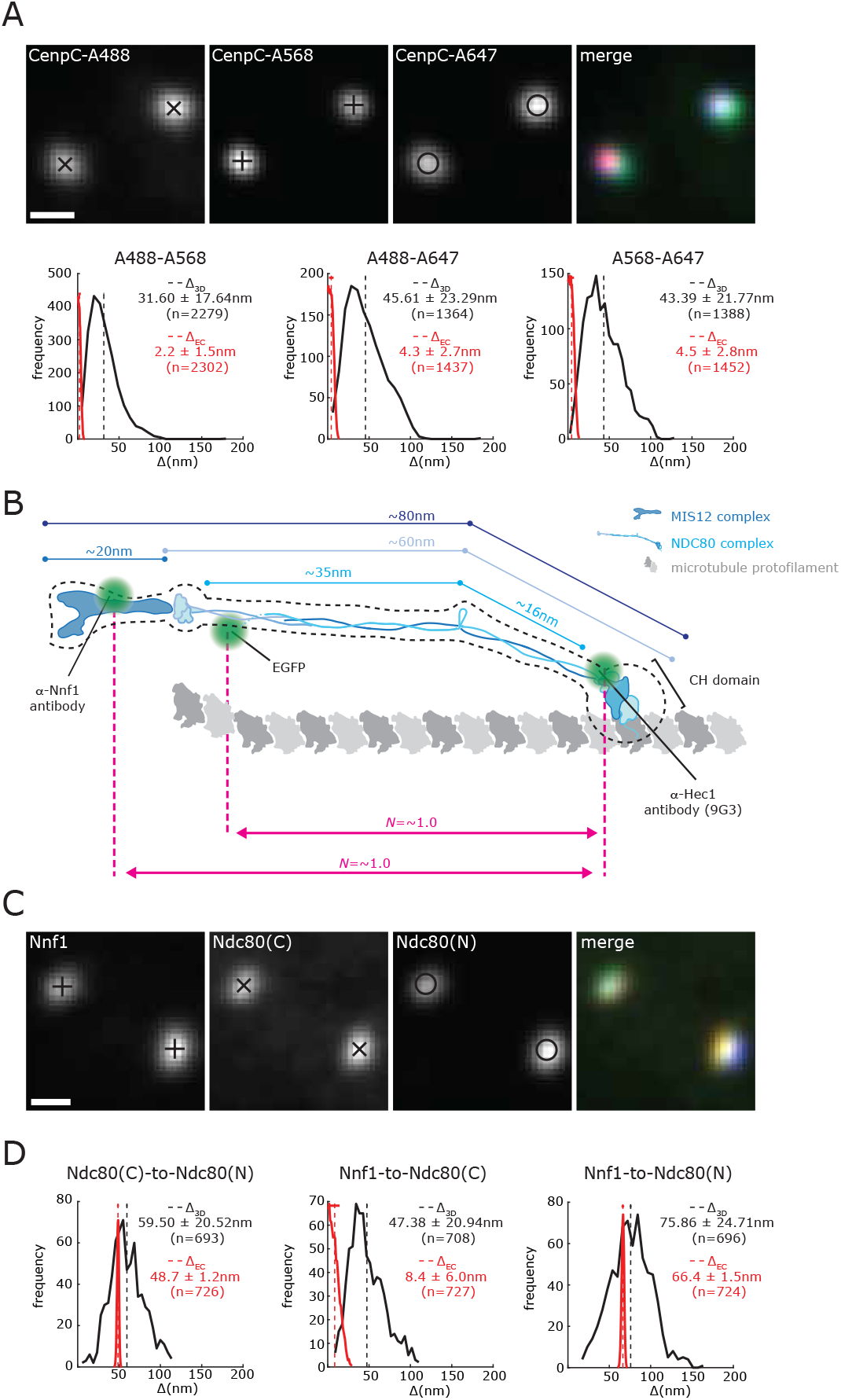
Δ_3D_ measurements overestimate the true Δ distance. (A) Kinetochores in RPE1 cells stained with anti-CenpC antibody and labelled with a mixture of Alexa488 (A488), Alexa568 (A568) and Alexa647 (A647)-conjugated secondary antibodies. Scale bar 500 nm. Histograms show Δ_3D_ and Δ_EC_ distances between the indicated fluorophores. Legend indicates distribution mean±sd. Vertical lines indicate means, bars on top dash show sd of estimated mean. (B) Schematic showing approximate organization and size of MIS12 complex (dark blue) and NDC80 complex (light blue) based on structural biology information available in literature. Dotted black line indicates the approximate shadow obtained from electron microscopy (EM) studies of MIS12 and NDC80 complexes. A single tubulin protofilament is shown in grey. Green dots indicate the approximate position of the antibodies and the GFP protein. Pink dotted lines and arrows indicate the distances used to calculate the nematic order (*N*) (see Supplemental Table 2). (C) Images of kinetochores in RPE1 Ndc80-EGFP cells stained with anti-Nnf1 and anti-Hec1(9G3) antibodies. Scale bar 500nm. (D) Histograms showing the Ndc80(C)-to-Ndc80(N), Nnf1-to-Ndc80(C) and Nnf1-to-Ndc80(N) Δ_3D_ and the Δ_EC_ distances measured in RPE1 Ndc80-EGFP cells. Legend indicates distribution mean±sd.

These data confirm that we can measure the 3D nanoscale distances between three different fluorophores within the human kinetochore with an accuracy on the scale of 2-4 nm. As a second test, the amino-terminal end of the Ndc80 subunit (of the NDC80 complex) was marked with the 9G3 monoclonal antibody that recognises the N-terminal globular domain of Hec1 (amino acids 200-215; De Luca et al., 2006; from here on referred to as Ndc80(N)) and the carboxy-terminal end with EGFP (Ndc80(C); Figure 1B, 1C). To label the Ndc80 C-terminal end we inserted EGFP at the endogenous locus by CRISPR/Cas9 gene editing in RPE1 cells (cell line MC191; see Materials and Methods and Supplemental Figure 2 for details). The Δ_EC_ distance from Ndc80(C)-to-Ndc80(N) was 48.7±1.2 nm (n=726; Figure 1D; Supplemental Table 1). Because there is substantial structural information describing the NDC80 complex *in vitro*, it has been used as a “molecular ruler” to validate two colour fluorescence localisation methods (Suzuki et al., 2018). Our Δ_EC_ measurements are close to the 51 nm distance estimated from negative stain electron microscopy (Wei et al., 2005, Huis In’t Veld et al., 2016; Figure 1B). As expected, the Δ_3D_ measurement is an overestimate (59.50±20.52 nm, n=693; Figure 1D; Supplemental Table 1), while the Δ_1D_ is typically an underestimate (33.37±14.48 nm, n=283; Supplemental Figure 1A and Supplemental Table 1) (Smith et al., 2016). We next measured the Δ_EC_ distance between the MIS12 complex subunit Nnf1 and Ndc80(N) and obtained a value of 66.4±1.5 nm (n=724, Figure 1D and Supplemental Table 1). This is similar to the distance measured in parental cells that did not have EGFP knocked-in to the *NDC80* locus (61.5±0.8 nm, n=1748; Supplemental Figure 1D; Supplemental Table 1). Again, these distances are very close to the predicted distance (~65 nm) that can be estimated from EM studies (Petrovic et al., 2010; Screpanti et al., 2011). Overall, we find that our algorithm gives tight confidence intervals similar to the accuracy determined for dual labels of the same protein (Supplemental Table 1), and that these data confirm the accuracy of our method and the importance of correcting intra-kinetochore distance measurements.

### The inner kinetochore is off-set from the outer kinetochore

Concurrent with measuring the distance between Ndc80(C) and Ndc80(N) we also determined an inner kinetochore position using the same CenpC antibody used in Figure 1A. We measured the distances from the amino- and carboxy-ends of Ndc80 to CenpC in a 3-fluorophore experiment and obtained a triangle with side lengths 43.5±2.3 nm (n=247), 81.9±2.9 nm (n=244) and 48.4±3.1 nm (n=247; Figure 2A, 2B; Supplemental Figure 1B; Supplemental Table 1). These 3 distances are non-compatible with collinearity (p=0.019, z= 2.07; Figure 3B; Supplemental Table 1). To further substantiate a lack of collinearity, we also measured the distances from Ndc80(N) and CenpC to Nnf1 in parental cells (Supplemental Figure 1C). These measurements also generate a significant triangle (p=6×10^−17^, z=8.28) with side lengths 34.9±0.6nm (n=1777), 85.8±0.8nm (n=1754) and 61.5±0.8nm (n=1748; Figure 2A, 2B; Supplemental Figure 1D; Supplemental Table 1). This contrasts to distances between Nnf1-Ndc80(C)-Ndc80(N) which are collinear to our accuracy (p=0.07, z=1.47, distances 48.7 ±1.2nm (n=726), 8.4±6.0nm (n=727) and 66.4±1.5nm (n=724); Figure 1D; Figure 2B; Supplemental Figure 1A; Supplemental Table 1). The CenpC position (inner kinetochore) must therefore be off-set (on average) from the axis defined by the Nnf1-Ndc80(C)-Ndc80(N) axis (outer kinetochore).

**Figure 2.**
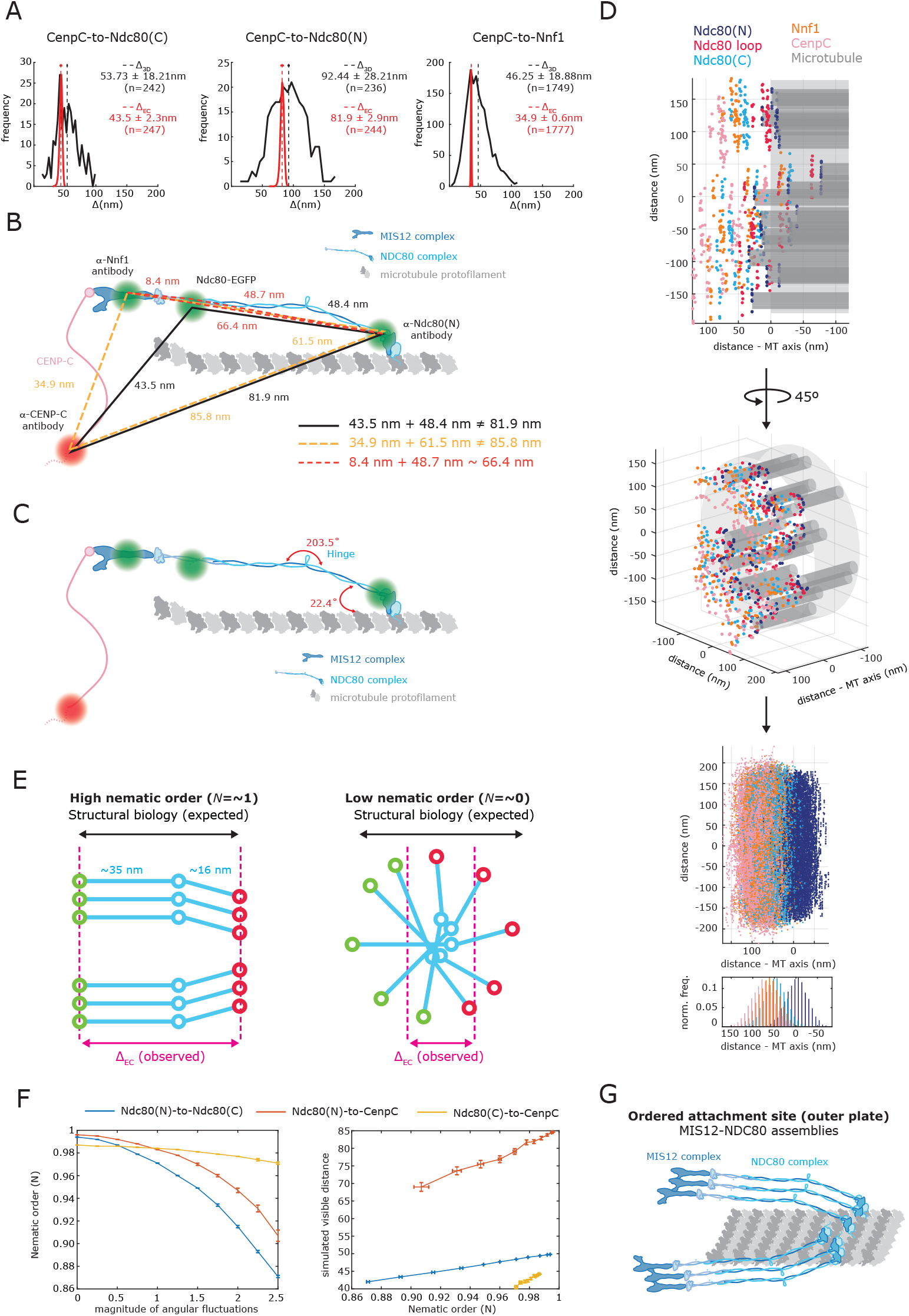
Simulations of the architecture of a bi-orientated kinetochore. (A) Histograms showing the CenpC-to-Ndc80(C) and CenpC-to-Ndc80(N) Δ_3D_ and the Δ_EC_ values measured in RPE1 Ndc80-EGFP cells and CenpC-to-Nnf1 Δ_3D_ and the Δ_EC_ distances in RPE1 cells. Legend indicates distribution mean±sd. (B) Schematic showing inferred architecture of CenpC-MIS12-NDC80. Green and red dots indicate the approximate position of antibodies and the GFP protein. Lines indicate measurements obtained in 3-fluorophore experiments between CenpC-Ndc80(C)-Ndc80(N) (black lines), CenpC-Nnf1-Ndc80(N) (orange dotted lines) and Nnf1-Ndc80(C)-Ndc80(N) (red dotted lines). (C) Schematic of CenpC-MIS12-NDC80 showing best fitted angles (in red) obtained from the simulations, see Supplemental Methods. (D) Simulated kinetochore organization. Top and middle panel show two perspectives of the same simulated kinetochore: each dot represents a marker on the designated molecule specified (by colour) in legend. Bottom panel shows an ensemble of 200 kinetochore simulations (aligned along their microtubule axis and relative to the mean of their Ndc80(N) markers) displaying the kinetochore layered structure along the microtubule (MT) axis direction. The bottom histogram shows the marker density along K-fibre for the above plot. See Supplemental Methods for simulation details. (E) Schematic representinghigh nematic order (*N*=~1) and low nematic order (*N*=~0) in the NDC80 complexes. Ndc80(C) and Ndc80(N) are shown as green and red circles, respectively. (F) Left panel: nematic order of the population of NDC80 complexes against the magnitude of the stochastic angular fluctuations (relative scale of the angular standard deviations compared to Supplemental Methods). Right panel: average distance between the indicated molecular markers against the nematic order. Simulations based on 200 kinetochores. (G) Schematic showing the inferred organization of MIS12-NDC80 complexes within a single microtubule attachment site in a bi-orientated kinetochore.

**Figure 3.**
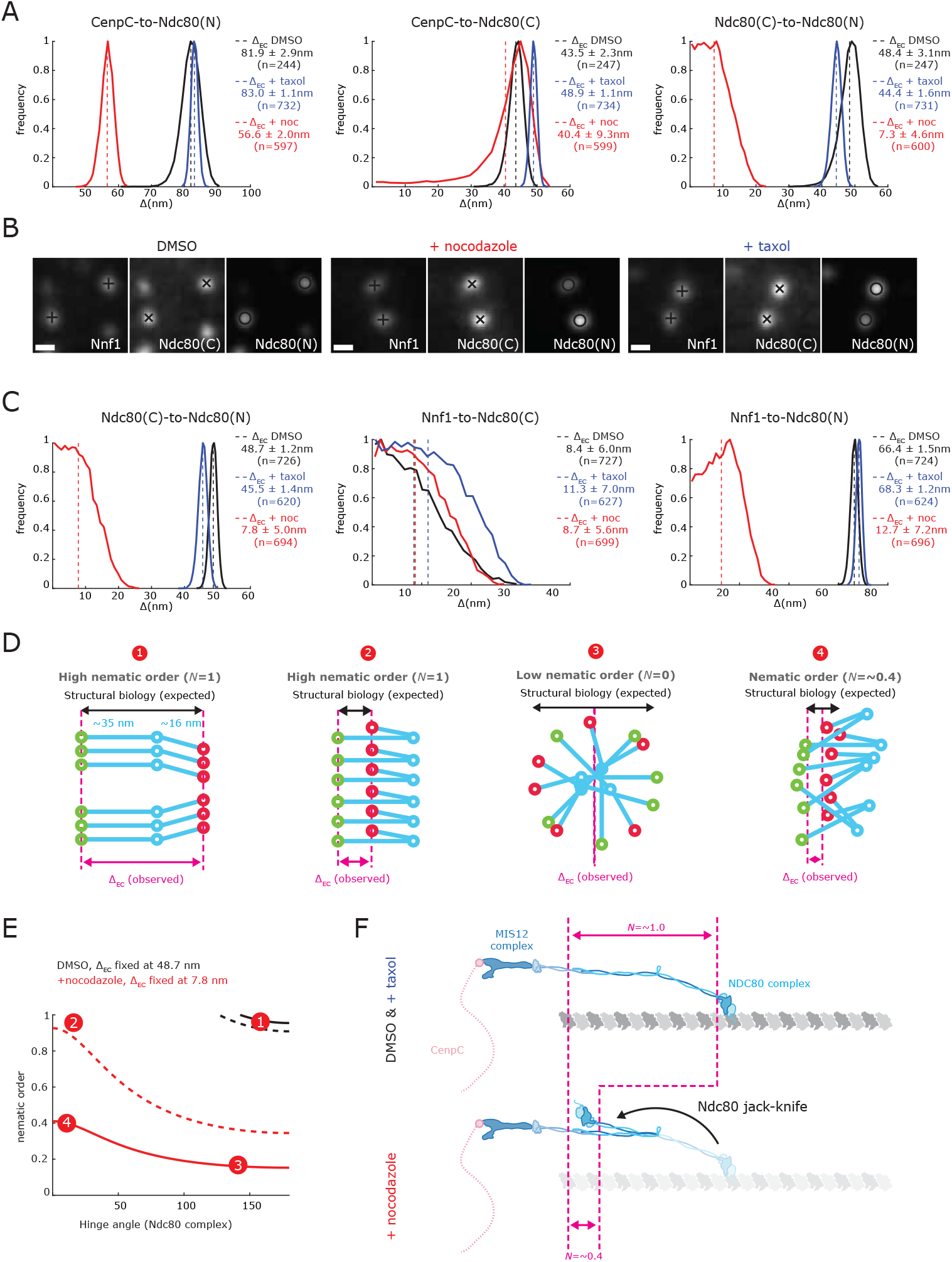
Loss of microtubule binding and stabilisation of microtubule dynamics induce different conformations of Ndc80. (A) Histograms showing the Δ_EC_ distance between CenpC-to-Ndc80(N), CenpC-to-Ndc80(C) and CenpC-to-Ndc80(N) in RPE1 Ndc80-EGFP cells treated with 3.3*μ*M nocodazole for 2hr, 1*μ*M taxol for 15 min or DMSO as control. Legend indicates distribution mean±sd. (B) Kinetochores in RPE1 Ndc80-EGFP cells stained with anti-Nnf1 and anti-Hec1(9G3) antibodies and treated with nocodazole, taxol and DMSO as control. Scale bars 500 nm. (C) Histograms showing the Δ_EC_ distance between Ndc80(C)-to-Ndc80(N), Nnf1-to-Ndc80(C) and Nnf1-to-Ndc80(N) in RPE1 Ndc80-EGFP cells in nocodazole, taxol and DMSO as control. Legend indicates distribution mean±sd. (D) Schematics showing 4 models of different structural and nematic organization of NDC80. High nematic order (*N*=1) is indicated when the observed Δ_EC_ coincides with the Ndc80(C)-to-Ndc80(N) distance expected from structural biology: Model 1 high Δ_EC_ with straight NDC80 conformation, Model 2 low Δ_EC_ with NDC80 jack-knife. However, low Δ_EC_ can also result from Model 3, a complete disorganized arrangement of straightened conformation NDC80 Model 3. Structural conformation and nematic order thus both contribute to reduced Δ_EC_, Model 4, *N*=0.4. (E) Data constrained nematic order and hinge angle for the NDC80 complex. For each hinge angle the nematic order is calculated that is consistent with the observed Δ_EC_, 48.7 nm (in DMSO) and 7.8 nm (in nocodazole) for the Ndc80(C)-to-Ndc80(N) linkage (see panel 3C). The four models of Figure 3D are shown: 1 and 2 are highly ordered arrangements (*N*=~1) of Ndc80 in control (1) and nocodazole (2)-treated cells. In nocodazole, the 7.8 nm distance between Ndc80(C) and Ndc80(N) can be explained by a completely disorganized NDC80 complex (position 3, *N*=<0.2) or a reduction in the hinge angle to almost 0° and nematic order of ~0.4 (position 4). Dashed lines show 95% confidence intervals. Hinge angles over 180° reflect around 180°. (F) Schematics representing the approximate arrangement of the MIS12 complex (dark blue) and NDC80 complex (light blue) in taxol- and nocodazole-treated cells with respect to control cells (faded colour schematics). CenpC is shown in pink and microtubule protofilament in grey. Pink dotted lines and arrows indicate the distances used to calculate the nematic order (*N*) (Supplemental Table 2).

### Visualisation of kinetochore ensemble organisation

To understand this lack of collinearity and interpret our distances in the context of known structural information, we built a computational simulation. It is important to note that each kinetochore contains multiple copies of each protein per microtubule, and multiple microtubule binding sites per kinetochore. Our experiments thus measure the ensemble average position of the tagged proteins, and not the distance between single molecules. Our simulations incorporate i) known structural biology on the NDC80 and MIS12 complexes, ii) information/measurements from electron microscopy data of K-fibre and kinetochore organisation and iii) measurements of kinetochore protein numbers per microtubule. We optimise unknown parameters by fitting the simulations to our measured triangle CenpC-Ndc80(C)-Ndc80(N). Crucially we fit the NDC80 complex hinge angle and the elevation of the NDC80 complex short arm (Ndc80(N)-to-Ndc80(hinge)) with the microtubule. Best fits are 203.5° and 22.4°, respectively (Figure 2C; See Supplemental Methods for details); hinge angles over 180° have been observed (Scarborough et al., 2019; Wang et al., 2008), whilst the latter is compatible with the range determined from EM and crystal structure studies (Ciferri et al., 2008; Wilson-Kubalek et al., 2008). Figure 2D (top and middle panels) shows a simulation of a single kinetochore bound to a K-fibre (grey) with the simulated positions of CenpC (pink), Ndc80(C) (light blue), Ndc80 loop (red), Ndc80(N) (blue) and Nnf1 (orange) single molecules. Averaging across multiple simulated kinetochores reveals a multi-layered structure with tight localisation and sequential ordering of kinetochore components along the K-fibre axis, and an extensive spread transverse to the axis suggestive of discs (Figure 2D, bottom panel). Our intra-fluorophore separation accuracy of 2-4 nm in our assay is substantially smaller than the kinetochore’s lateral and transverse dimensions. Our simulations indicate that moderate levels of flexibility of the Ndc80 microtubule attachment angle and orientation can produce a significant triangle between the average positions of CenpC, Nnf1/Ndc80(C) and Ndc80(N). In addition, the broad transverse dimension allows even small re-distributions of the CenpC population to also give rise to a significant triangle. Thus, changes in the distribution of the CenpC marker - potentially due to CenpC’s inherent flexibility - are sufficient to explain the lack of co-linearity measured *in vivo*.

The layered structure along the K-fibre axis is a result of high molecular alignment, *i.e*. the NDC80 complexes have high nematic order along the K-fibre axis reminiscent of liquid crystals. This high alignment allows comparison between structural data and our *in vivo* distance measurements. Specifically, under high order the distances should match, whilst as nematic order decreases the measured distance of the ensemble decreases (Figure 2E). The angular degrees of freedom, such as the NDC80 hinge (Supplemental Methods) are in effect highly constrained to produce high alignment of the NDC80 complexes. We investigated how increasing the fluctuations in the hinge angle and Ndc80 orientation decreased the nematic order, Figure 2F (left panel) and correspondingly decreased the average Δ_EC_ distance Figure 2F (right panel). Thus, the high nematic order in the outer plate is essential for comparison with structural data.

We can quantify this alignment order from our vivo measurements with an order statistic *N* (see Supplemental Table 2), where *N*=~0 indicates no alignment (low nematic order), and N=~1 is perfect parallel alignment of vectors (high nematic order; Figure 2E). We find that for both the Ndc80(C)-to-Ndc80(N) and Nnf1-to-Ndc80(N) linkages *N* = ~1 (Figure 1B; Supplemental Table 2), assuming the Ndc80 hinge angle is 203.5°. This indicates that in the microtubule-attached state the MIS12-NDC80 complex assemblies are highly ordered and that Δ_EC_ distances reflect the underlying molecular organisation (Figure 2G) as implicitly assumed in using the NDC80 complex as a “molecular ruler” (Suzuki et al., 2018). Thus, our simulations and data indicate that at this resolution, the kinetochore is organised along an internal axis with high nematic order, particularly in the outer plate (Supplemental Table 2).

### NDC80 complex jack-knifes on unattached, but not tensionless kinetochores

We next investigated how the structural organisation and geometry of the kinetochore responds to a loss of attachment or tension. To do this, we treated the RPE1 cell lines with either i) 3.3 *μ*M nocodazole for 2 hr and confirmed by tubulin staining that all kinetochore-microtubule populations were eliminated and that the 3D K-K distance (CenpC to CenpC) was reduced (0.93±0.005 *μ*M, n=2537, mean±SEM), or ii) 1 *μ*M taxol for 15 min which reduces K-K distance to nearly rest length (0.96±0.004 *μ*m, n=1884, mean±SEM), but leaves the majority of kinetochores associated with microtubules and aligned on the metaphase plate (Supplemental Figure 3A). We next checked how our 1 *μ*M taxol treatment affects microtubule dynamics by photoactivating PA GFP-alpha-tubulin adjacent to the metaphase plate and then measuring its movement to the pole (poleward microtubule flux) and the dissipation of the signal over time (plus-end turnover). Clearly, treatment with 1 *μ*M taxol abolishes plus-end turnover as the tubulin signal is stable for 120 s, while the signal in DMSO treated cells exponentially decays as previously reported (Amaro et al., 2010); Supplemental Figure 3B, 3C). The flux rate was also reduced from 0.76±0.36 *μ*m min^−1^ in DMSO treated cells to 0.04±0.25 *μ*m min^−1^ in taxol-treated cells (p=3.5×10^−8^; Supplemental Figure 3B, 3C). Thus 1 *μ*M taxol largely eliminates microtubule dynamics consistent with the loss of tension.

In a first experiment, we measured the geometry of the CenpC-MIS12-NDC80 assembly: the distances from CenpC to Ndc80(C) and Nnf1 were largely unchanged in nocodazole and taxol (Figure 3A; Supplemental Figure 4A, 4B; Supplemental Table 1). This is consistent with previous work reporting that the inner kinetochore is a stiff structure (Smith et al., 2016; Suzuki et al., 2018). However, the CenpC-to-Ndc80(N) distance was reduced by 25.3±3.5 nm following treatment with nocodazole (in RPE1 Ndc89-EGFP cells, Figure 3A; similar to the reduction of 30.1±3.3 nm observed in parental cells, Supplemental Figure 4B; Supplemental Table 1). Consistent with the reduced CenpC-to-Ndc80(N) linkage, the Ndc80(C)-to-Ndc80(N) distance decreased by 41.1±5.5 nm (85%) in unattached kinetochores (Figure 3A). We also measured the distances between Nnf1, Ndc80(C) and Ndc80(N) (Figure 3B). This triangulation confirms there is a substantial reduction of 40.9±5.1 nm (84%) in the Ndc80(C)-to-Ndc80(N) linkage under nocodazole treatment (Figure 3C; Supplemental Table 1). This triangulation also demonstrates that the distance between Nnf1 and Ndc80(C) remains unchanged in nocodazole (Figure 3C; Supplemental Table 1), and surprisingly, in taxol the Δ_EC_ between Ndc80(C) and Ndc80(N) remained the same as in control (Figure 3A and 3C; Supplemental Table 1). This suggests that the Ndc80 N-terminus does not move inwards following taxol treatment and the NDC80 complex remains in its straight configuration with a high nematic order, *N*=~1 (Figure 3F and Supplemental Table 2). These data provide evidence that the kinetochore responds differentially to loss of attachment or tension.

We propose that under loss of microtubule attachment the NDC80 complex has undergone an intra-molecular jack-knife movement bringing the N- and C-termini into close proximity (Δ expected = ~19 nm based on structural data, see Supplemental Table 2; Figure 3D, Model 1 versus Model 2). However, given the ensemble nature of the kinetochore the data could also be explained by a loss of nematic order without any change in NDC80 complex organisation (*e.g*. Figure 3D, Model 3 shows a total loss of order). Only for rigid molecules is the nematic order the ratio of the observed to structural distance; for molecules with internal flexibility a change in the observed distance can be a consequence of both a change in the molecule flexing and in alignment. We therefore examined how the data can be reproduced through a combined change in the NDC80 hinge angle and nematic order (Figure 3E). This demonstrates that if the Ndc80 jack-knifes into a ridged closed conformation, a reduction in the nematic order to 0.4 will be needed for Δ_EC_ to equal the measured 7.8 nm (Model 4 in Figure 3D and 3E). Consistently, the nematic order for Ndc80(C)-to-Ndc80(N) is 0.4 in nocodazole-treated cells (Figure 3F; Supplemental Table 2). On the other hand, if there were no jack-knife in the NDC80 complex (it remains straight and rigid) then the order would need to decrease to <0.2 (i.e. almost complete loss of order) to explain the measured 7.8 nm distance (Figure 3E, Model 3). We do not, however, favour this idea because the distance from Nnf1-to-Ndc80(C) does not change under nocodazole treatment (8.4±6 nm *vs*. 8.7±5.6 nm; Figure 3C); *i.e*. there is no evidence for a loss of order in the MIS12 complex to NDC80 complex junction. Taken together, these data show that the NDC80 complex N-terminus jack-knifes in the absence of microtubule binding.

### Ndc80 jack-knifing marks spindle checkpoint active kinetochores

Do these conformational changes relate to checkpoint signalling from the kinetochore? For these experiments we used RPE1 cells in which Mad2 was labelled at its endogenous locus with Venus (Collin et al., 2013). As expected nocodazole treatment (unattached kinetochores) leads to an increase in Venus-Mad2 and Bub1 kinetochore binding on most kinetochores while after taxol treatment the average levels of these proteins were similar (Figure 4A, 4B; Supplemental Figure 5A). However, in taxol and DMSO there were multiple outliers (Figure 4B) suggesting that there is a Mad2 positive subpopulation as previously reported (Magidson et al., 2016). In addition, we found that treatment with 1 *μ*M taxol prevents cells in metaphase from exiting mitosis unless 1 *μ*M reversine is added to the media (Figure 4C; Supplemental Figure 5B). These experiments indicate that the spindle checkpoint is active (or not silenced) following treatment with 1 *μ*M taxol and suggest that the few Mad2-signalling kinetochores might be responsible. To understand how the Ndc80 jack-knife relates to Mad2 recruitment we separated our kinetochore spots into a VenusMad2 positive (Mad2^+^) and negative (Mad2^−^) population, and then determined the distance from CenpC-to-Ndc80(N) (Figure 4D). The CenpC-to-Ndc80(N) distances were significantly different in these two populations (p=2.4×10^−5^), with the distance in the Mad2^+^ population (66.2±3.7 nm) nearly identical to that of nocodazole treated cells (all Mad2^+^, 58.3±1.3 nm, p=0.044). Further, the difference of the CenpC-to-Ndc80(N) of 16.2±3.8 nm between Mad2^−^ and Mad2^+^ in DMSO (Figure 4D; Supplemental Table 1) is consistent with the Ndc80 jack-knifing under lack of attachment in Mad2^+^ (compare to the 25.3±3.5 nm under nocodazole, p=0.039) suggesting that there is a high correlation between Mad2^+^/Mad2^−^ and Ndc80 being in the jack-knife or straight conformations (nocodazole versus DMSO, respectively). Under taxol, we could also detect an inward movement of 11.5±5.1 nm of the Ndc80(N) in the Mad2^+^ versus Mad2^−^ subpopulation (Figure 4D). These correlations are imperfect because our Mad2^+^/Mad2^−^ populations may not be optimal due to noise, and there is likely a contribution from a partial jack-knifing state as error correction proceeds.

**Figure 4.**
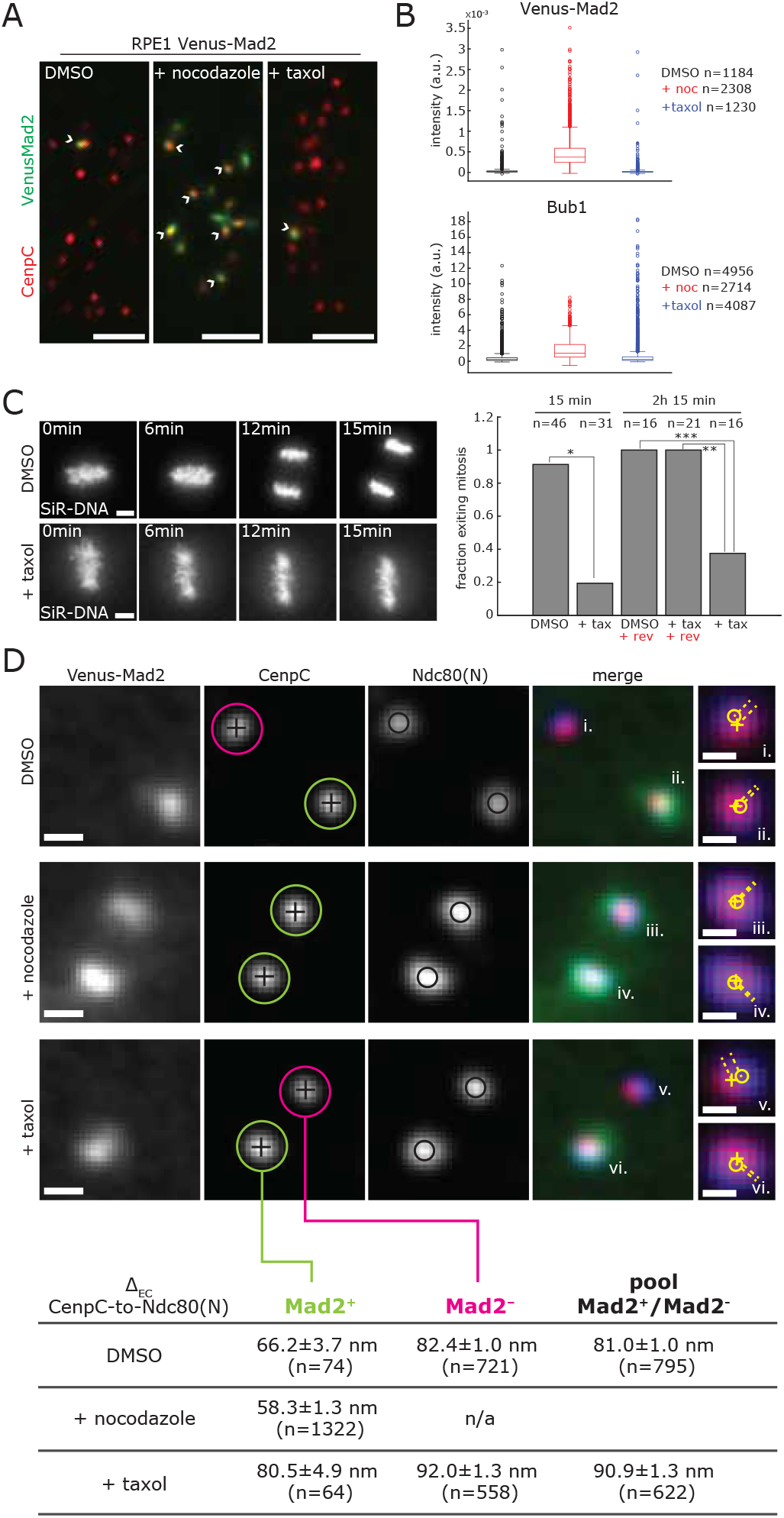
Activation of the Spindle Assembly Checkpoint correlates with NDC80 conformational change. (A) Images of RPE1 Venus-Mad2 cells stained with anti-CenpC antibody and treated with 3.3 *μ*M nocodazole for 2 hr, 1 *μ*M taxol for 15 min or DMSO as control. White arrowheads indicate Mad2 signalling kinetochores. Scale bars 3 *μ*m. (B) Box and whiskers plots displaying Venus-Mad2 and Bub1 intensity at kinetochores treated with nocodazole (+noc), taxol and DMSO. Venus-Mad2 signal is background-subtracted. Bub1 signal is background-subtracted and normalised to CenpC signal (also background corrected). (C) Images of RPE1 cells treated with 1 *μ*M taxol and DMSO for 15 min. To visualize DNA, cells were stained with SiR-DNA. Scale bars 5 *μ*m. Bar chart on the right shows the fraction of cells exiting mitosis within the indicated imaging time, in DMSO, taxol (+tax) and upon addition of 1 *μ*M reversine (+rev) (Supplemental Figure 5B; see Material and Methods for details). Fisher’s exact test indicates the differences are significant with 99% confidence interval: (*) p=7.7×10^−12^; (**) p=2.3×10^−5^; (***) p= 1.2×10^−4^. (D) Example images of kinetochores in RPE1 Venus-Mad2 cells stained with anti-CenpC and anti-Hec1(9G3) antibodies, and treated with nocodazole, taxol and DMSO. Scale bars 500nm. Table at the bottom displays the Δ_EC_ (mean±sd.) between CenpC and Ndc80(N) in Venus-Mad2 positive (green circle), Venus-Mad2 negative (pink circles) kinetochores and pool samples, in the 3 conditions. Insets (i. to vi.) show the distances (yellow dotted lines) between CenpC (+) and Ndc80 (O) in the indicated kinetochores on the XY plane. Scale bars in insets 250nm.

Importantly, these data indicate that we can detect the inward movement of Ndc80(N) at Mad2^+^ kinetochores in control metaphase cells, and not just under nocodazole treatment. Together these data clearly demonstrate that checkpoint activation (in terms of Mad2 recruitment) occurs at kinetochores that are in the unattached conformation (Ndc80 jack-knifed). Moreover, our data show how most kinetochores in 1 *μ*M taxol have a low K-K distance without triggering the recruitment of Mad2.

### Mad2 can occupy two distinct positions within the kinetochore

We next sought to establish where in the kinetochore Mad2 is recruited. While Bub1 is a *bona fide* kinetochore receptor for human Mad1:Mad2 complexes (Zhang et al., 2017), the RZZ complex, is also implicated (Kops et al., 2005) and may even operate as a separate receptor (Currie et al., 2018; Silio et al., 2015). To read out the position of Bub1 we used an antibody that recognises the first 300 amino acids (referred to as Bub1 (N), Figure 5B) and found it positioned 58.9 ± 1.1 nm (n=2232) from CenpC and therefore 26.9±1.4 nm to the inside of the Ndc80 head domain (Figure 5C, Supplemental Table 1). Using antibodies that recognise the carboxy-terminus of Rod (referred to as Rod(C)) and Zwilch (Figure 5A, 5B), we found that while Rod(C) is positioned 14.4±2.1nm inside Ndc80(N), Zwilch is 10.5±1.3nm to the outside (Figure 5B, 5C, Supplemental Table 1). These results indicate that most of the RZZ complex is located outside the Ndc80 head domain, and thus is spatially separated from Bub1 (Figure 6F, DMSO). We can estimate the distance between Zwilch and Rod by subtracting the CenpC-to-Rod(C) from CenpC-to-Zwilch distances giving 24.9±2.1 nm (Supplemental Table 1). This distance is similar to that determined from the cryo-EM structure (Mosalaganti et al., 2017) and indicates high nematic order (*N*=~0.8; Supplemental Table 2). The high nematic order allows us to use the ‘molecular ruler’ to organise the outer kinetochore/corona. This data also suggests that, at least in end-on attached kinetochores, the RZZ complex is not forming a head-to-tail dimer as suggested by cryo-EM (Mosalaganti et al., 2017; which would have resulted in Rod(C) and Zwilch signals being coincident). These data thus provide the first 3D mapping of the key checkpoint protein platforms (Bub1 and RZZ) relative to the major microtubule attachment factor (NDC80 complex) in the human kinetochore.

**Figure 5:**
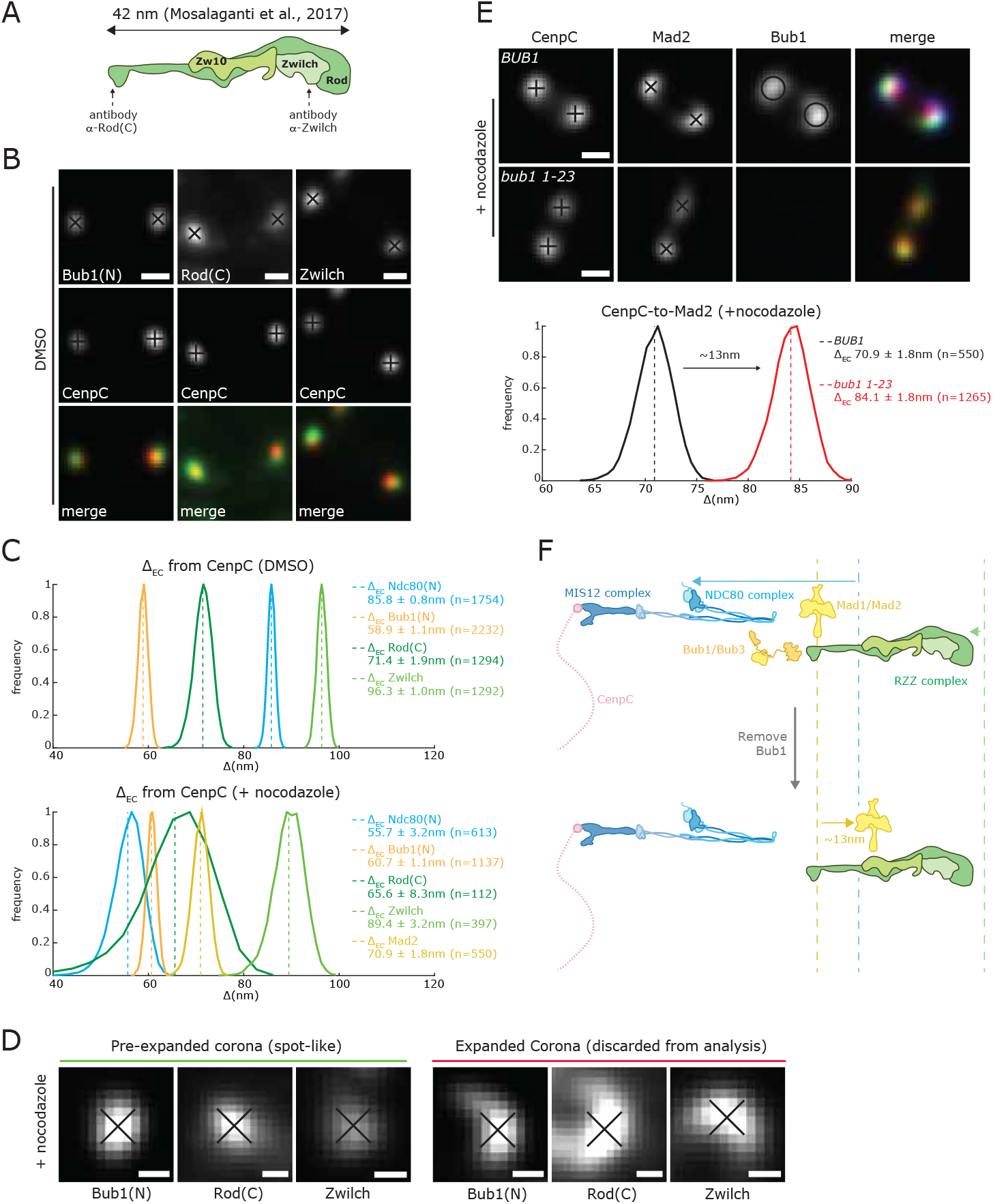
Mad2 binds different kinetochore sites upon activation of the Spindle Assembly Checkpoint. (A) Schematics showing the RZZ complex size and the approximate binding positions of Rod(C) and Zwilch antibodies used in this study. (B) Kinetochore pairs stained with anti-CenpC antibody in combination with either anti-Bub1, anti-Rod or anti-Zwilch antibodies in control cells (DMSO treated). Scale bars 500nm. (C) Histograms showing the Δ_EC_ between CenpC and Ndc80(N), Bub1, Rod and Zwilch in DMSO (top) and nocodazole (bottom) treated cells. Mad2 position shown for nocodazole only. Legend indicates distribution mean±sd. (D) Images of Bub1, Rod and Zwilch kinetochores treated with 3.3*μ*M nocodazole for 2 hr showing examples of spot-like and expanded kinetochores. Scale bars 250nm. (E) Images of kinetochores in parental RPE1 and RPE1 *bub1 1-23* cells stained with anti-CenpC, anti-Bub1 and anti-Mad2 antibodies and treated with 3.3 *μ*M nocodazole for 2hr. Scale bars 500 nm. Lower histograms show the Δ_EC_ between CenpC and Mad2 in parental and RPE1 *bub1 1-23* cells in nocodazole. Legend indicates distribution mean±sd. (F) Schematics representing the approximate arrangements of the MIS12 complex (dark blue), NDC80 complex (light blue), Bub1/Bub3 (orange) and RZZ complex (green) in parental RPE1 (top schematic) and in RPE1 *bub1 1-23* cells (bottom schematic) upon nocodazole treatment. CenpC is shown in pink. Dotted lines represent the position of the indicated complexes in control cells (DMSO). Arrows indicate the change in position between different conditions.

We mapped the positions of Mad2 and Bub1 utilising the RZZ molecular ruler. To measure the Mad2 position within kinetochores, cells were treated with nocodazole to depolymerize microtubules and trigger recruitment of Mad2. Because the corona/outer kinetochore begins expanding into crescent-shaped structures under these (unattached) conditions we limited our analysis to non-expanded kinetochores in order to have accurate positional information (Figure 5D). In the absence of microtubules, the position of Bub1, Rod and Zwilch (RZZ complex) remained largely unchanged (Figure 5C, 5F). This clearly shows that kinetochores do not simply collapse when unattached. Instead, there are well defined structural rearrangements.

At unattached kinetochores, Mad2 was located 70.9±1.8 nm from CenpC, which is positioned outside of Bub1 and close to the position of Rod(C) (Figure 5C, Supplemental Table 1). We hypothesised that the lack of co-location with Bub1 may reflect Mad1:Mad2 binding to both Bub1 and a second receptor proximal to the RZZ complex (Silio et al., 2015); the average position moving to the outside of Bub1 (note that our measurements are the average position of a protein within a kinetochore and across multiple kinetochores). To test this idea, we measured the position of Mad2 in cells carrying a homozygous *bub1 1-23* mutation. In these cells Bub1 levels are reduced to almost undetectable levels but Mad2 can still bind kinetochores and signal the Spindle Assembly Checkpoint in cells treated with 1 *μ*M taxol (Supplemental Figure 6A, 6B; (Currie et al., 2018; Zhang et al., 2019)). In the presence of nocodazole, the Mad2 signal was now repositioned 13.2±2.5 nm further outwards (Figure 5E and 5F). This data is consistent with the model that there is a Bub1-independent kinetochore-receptor for Mad1:Mad2 that is dependent on the RZZ complex (Silio et al., 2015).

### Kinetochores adopt a unique conformation following loss of tension

The absence of a conformational change in NDC80 following taxol treatment (Figure 3F) raised the question as to whether loss of tension is detected at all. We first checked the position of Bub1 and RZZ subunits in taxol-treated cells and also found no changes (Figure 6A and 6B). We then turned to Knl1, the third component of the KMN network (Cheeseman et al., 2006). Knl1 is a largely disordered protein that binds to the MIS12 complex through the carboxy-terminus with the remaining protein comprising multiple MELT sequences that operate as phospho-dependent binding sites for the Bub3-Bub1 checkpoint complexes (Figure 6C; (London et al., 2012); (Shepperd et al., 2012; Yamagishi et al., 2012). The extreme amino-terminal end of Knl1 contains a microtubule-binding site and a docking site for protein phosphatase 1 (PP1) (Figure 6C). The amino terminus of Knl1 (marked with a phospho-specific antibody that recognises serine 24, referred to as pKnl1(S24)) and the second MELT (amino acids 300-350, referred to as Knl1(MELT2)) (Figure 6C) were both positioned in close proximity to Bub1 (12.2±10.6 and 6.5±1.5 nm, respectively; Figure 6B, 6F, DMSO). These data suggest that the unstructured region of Knl1 (Figure 6C) which has a predicted path length of ~500 nm must be “wrapped up” and occupy the space between the Ndc80 head domain and the CCAN (CenpC here) (Figure 6F, DMSO). We next checked the position of the amino-terminus of Knl1 (pKnl1(S24)) following taxol treatment (Figure 6D) and, remarkably, found that it moved outwards by 68.5±10.7 nm (Figure 6B, 6F). There was minimal movement of the second MELT, indicating that the bulk of the MELT array is unchanged (Figure 6B, 6E and 6F; Supplemental Table 1). This is consistent with our observation that Bub1 position – which is a proxy for the MELTs – does not move. Thus, the Knl1 amino-terminus moves from a position internal to MELT2 to a new position outside. The derived distance from Ndc80(N) is now 28.8±2.7 nm (Figure 6F, taxol). The predicted length of the disordered first 300 amino acids is ~64 nm, thus this region of Knl1 appears to switch into an almost straight configuration. We did not detect an increase in phosphorylation of Knl1 (S24), (Figure 6D), which is a substrate for the AuroraB kinase (Welburn et al., 2010). This suggests that these kinetochores remain attached and are not undergoing error-correction (see discussion). As expected, following treatment with nocodazole there was an increase in Knl1(S24) as these kinetochores are detached (Figure 6D). Because such kinetochores also loose tension we checked the position of Knl1 in nocodazole (Figure 6D, 6E). Knl1 is again more extended with the amino-terminal end now 94.8±3.6 nm (n=421) from CenpC (compared to 115.2±2.3 nm, n=492 in taxol; Figure 6D, 6F; Supplemental Table 1). We also detected an outward movement (17.1±2.4 nm) of the MELT2 position (Figure 6E and 6F, nocodazole). These data provide evidence that the physical re-organisation of Knl1 responds to the loss of tension.

**Figure 6.**
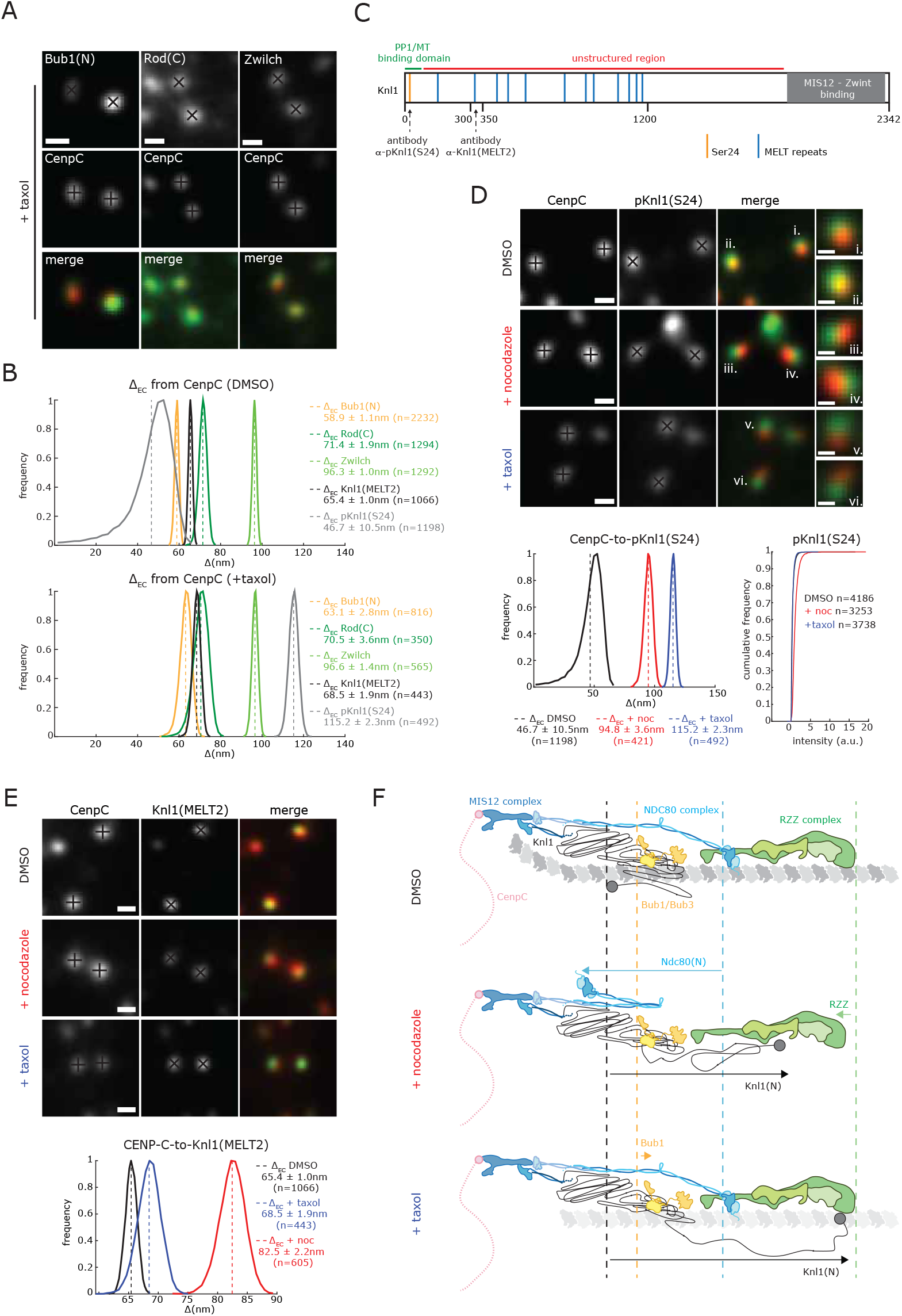
Knl1 1-300 unravels upon loss of kinetochore tension. (A) Kinetochore pairs stained with Bub1, Rod and Zwilch antibodies and treated with 1 *μ*M taxol for 15 min. Scale bars 500 nm. (B) Histograms showing the Δ_EC_ of Bub1, Rod, Zwilch, Knl1-MELT2 and pKnl1(S24) from CenpC in DMSO (top) and taxol (bottom). Legend indicates distribution mean±sd. (C) Schematic map of Knl1 where the positions of Serine24 (Ser24, orange bar), MELT repeats (light blue bars) and MIS12-Zwint binding domain (grey box) are shown. Green and red lines indicate the PP1/microtubule (MT) binding site and the unstructured region, respectively. Arrows indicate the binding sites of pKnl1(Ser24) and Knl1(MELT2) antibodies used in this study. (D) Kinetochores stained with anti-CenpC and anti-pKnl1(S24) antibodies and treated with 3.3 *μ*M nocodazole for 2 hr, 1 *μ*M taxol for 15 min or DMSO as control. Scale bars 500nm. Insets show enlargements for the indicated kinetochores. Scale bars 250nm. Histograms at the bottom show the Δ_EC_ between CenpC and pKnl1(S24) in DMSO, nocodazole (+noc) and taxol treated cells. Legend indicates distribution mean±sd. Cumulative frequency plots display the pKnl1 (S24) intensity in DMSO, nocodazole (+noc) and taxol treated cells. pKnl1(S24) signal was background subtracted and normalised to CenpC signal (also background corrected). (E) Kinetochores stained with anti-CenpC and Knl1(MELT2) antibodies and treated with 3.3*μ*M nocodazole for 2 hr, 1 *μ*M taxol for 15 min or DMSO as control. Scale bars 500nm. Histograms at the bottom show the Δ_EC_ between CenpC and Knl1 in DMSO, nocodazole (+noc) and taxol treated cells. Legend indicates distribution mean±sd. (F) Schematics representing the approximate arrangements of the MIS12 complex (dark blue), NDC80 complex (light blue), Bub1/Bub3 (orange), RZZ complex (green), Mad1/Mad2 (in yellow) and Knl1 (black) in RPE1 cells in DMSO, nocodazole and taxol. CenpC is shown in pink, microtubule protofilament in grey. Dotted lines represent the position of the indicated complexes in control cells. Arrows indicate the change in position between different conditions.

## Discussion

This work provides an initial architectural map of the human kinetochore (in diploid non-transformed cells), quantifying the relative position (accuracy 1-10 nm) and movement of major complexes and subunits. Our distances are corrected for Euclidean distance inflation and interpreted within the context of a kinetochore ensemble of molecules and available structural data. There is also an averaging over functional states. This was evident in that we can separate out Mad2^+^ and Mad2^−^ populations of kinetochores with significantly different intramolecular distances. Using a computational simulation, we generated a 3D visualisation of a human kinetochore which produces plate-like structures for the outer (MIS12-NDC80) and inner (CenpC) kinetochore components, reminiscent of that observed in electron micrographs (Brinkley et al., 1992). These data also demonstrate how the inner and outer plates in individual kinetochores are shifted relative to each other in the direction perpendicular to the microtubule axis by intrinsic stochasticity in the positioning and conformation of NDC80 complexes. We propose here that the outer kinetochore can be thought of as having properties of a liquid-crystal system, whereby the molecules align along an outer kinetochore axis, presumably parallel to the K-fiber axis. Within the outer plate the NDC80 complex, and the more distal RZZ complex, must have a high nematic order (otherwise the ensemble average distances would not be consistent with distances from structural biology). It thus follows that our methods allow structural insights from cryo-EM and X-ray crystallography to be assessed within an *in vivo* context. They also provide a framework for understanding the higher order ensemble organisation of the kinetochore.

At the start of mitosis kinetochores are in an unattached state and our data reveals how the Ndc80 head domains will be pivoted at the loop sequence in a folded back position (Figure 7, step 1). Recent *in vitro* experiments have confirmed that the NDC80 complex can indeed adopt such a state, and suggest that this is an auto-inhibited configuration that is relieved on microtubule binding (Scarborough et al., 2019). The folded back state is tightly correlated with the recruitment of Mad2 to kinetochores and activation of the spindle checkpoint. As end-on attachments form the NDC80 complexes will straighten out and align. During this time, the Mps1 kinase will be displaced from the Ndc80 calponin homology domain, competed off by the microtubule interaction (Figure 7, step 2; (Hiruma et al., 2015; Ji et al., 2015)). One idea is that the auto-inhibited Ndc80 conformation gates the recruitment of Mad1:Mad2, perhaps by favouring Mps1 binding. This model is different from that presented in budding yeast by (Aravamudhan et al., 2015) which proposes that attachment separates Mps1 from the Knl1 substrate, thus silencing the checkpoint. However, in bi-orientated human kinetochores the Ndc80 head domains and Knl1 MELTs (marked by average position of Bub1) are only 26.9±1.4 apart. Nevertheless, conformational changes in Ndc80 appear to be a conserved feature of the mechanisms that monitor microtubule attachment.

**Figure 7.**
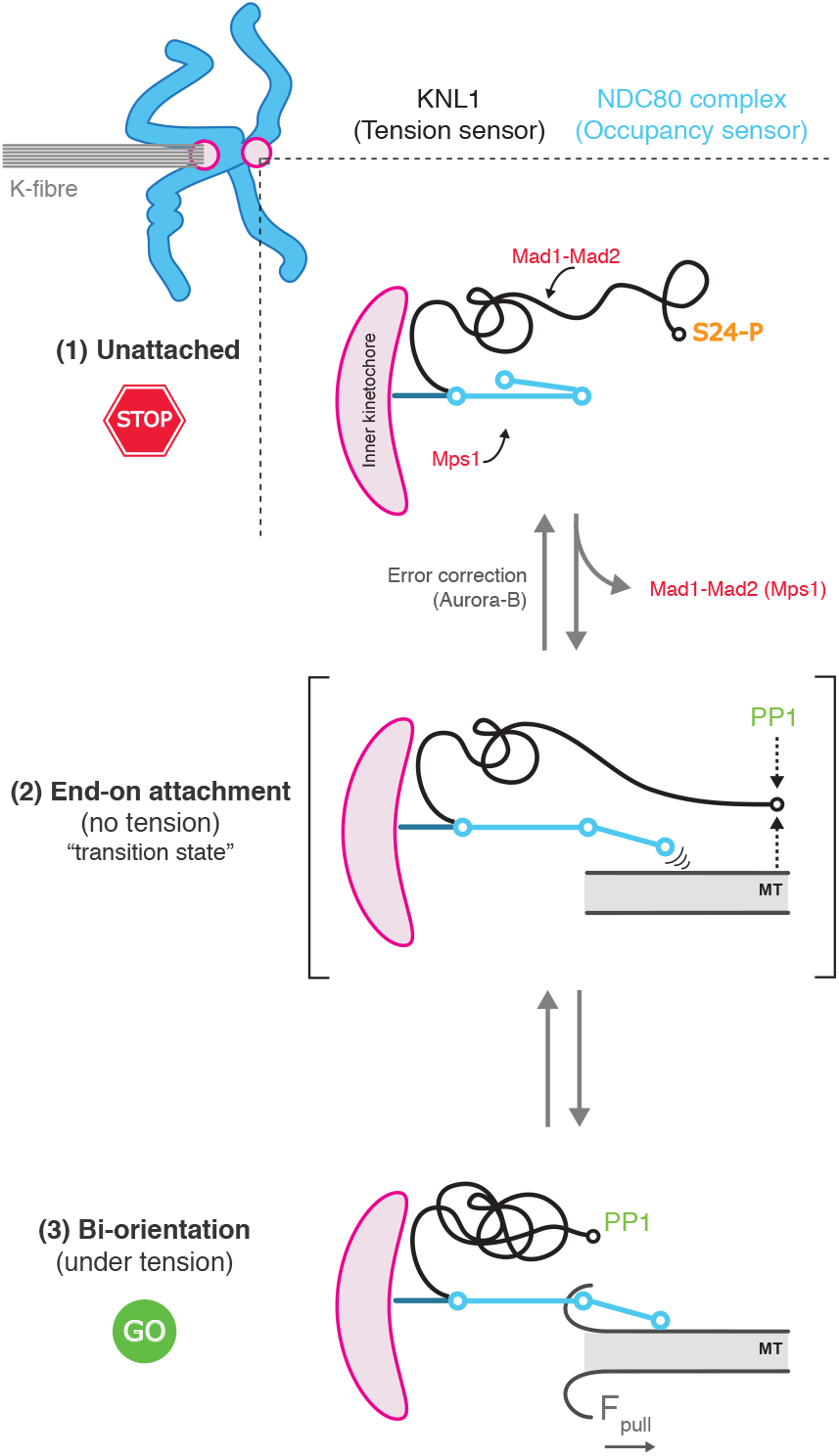
Model for the integration of tension and occupancy sensors with checkpoint and error correction mechanisms. At unattached kinetochores (top panel, 1) Ndc80 (light blue) is in auto-inhibited state and Knl1 (black) 1-300 aa is in extended conformation. Aurora B kinase activity dominates and kinetochore substrates (i.e. Ndc80, Knl1) are phosphorylated and in a low-affinity microtubule-binding state. Checkpoint proteins (red) are also bound and a “wait anaphase” signal is generated. When an end-on attachment forms (middle panel, 2) - but no tension generated (for example monotelic attachments where one sister is attached and the other unattached) - the checkpoint signalling is silenced on the attached sister concomitantly with straightening of Ndc80 which binds to the microtubule lattice (grey). We propose that recruitment of PP1 is slowed because binding of PP1 to the Knl1 amino-terminus is competitive with microtubule binding but sufficient to keep Knl1(S24) dephosphorylated. This phosphatase activity is also spatially separated from outer kinetochore substrates (*i.e*. Ndc80). As a result, microtubule binding by the kinetochore remains in a weakened meta-stable state. As tension is generated (bottom panel, 3) the Knl1 amino-terminus ravels and PP1 now fully binds, perhaps as microtubule binding is unfavourable. This leads to outer kinetochore dephosphorylation (counteracting Aurora B) and increased microtubule binding affinity. This model can also explain error correction following loss of tension at a bi-orientated kinetochore, which would switch the system into the transient (middle panel) state (see discussion for details).

Intriguingly, when kinetochores are attached - but no tension is generated (taxol state in this study) - we find that the first 300 amino acids of Knl1 are unravelled, while NDC80 does not undergo jack-knifing and an increase in Mad2-kinetochore binding cannot be detected (Figure 4D). What drives cycles of unravelling-ravelling in the first 300 amino acids of Knl1 remains unknown. This supports the idea that loss of tension alone is not relayed to the spindle checkpoint (Magidson et al., 2016), although we cannot rule out weak checkpoint signals (without detectable Mad2 binding). Instead, we propose that Knl1 unravelling is part of a kinetochore “tension sensor” that is involved in error correction and the force-dependent stabilisation of microtubule attachments (See model in Figure 7 for details). The principle idea is that the outward movement of the Knl1 amino-terminus (when tension is lost) would spatially separate the bound phosphatase (PP1) from substrates in the outer kinetochore (*e.g*. Ndc80, Ska). This would enable AuroraB to dominate and attachment stability to be weakened (ultimately leading to error correction), or correspondingly attachment stabilisation when sister kinetochores come under tension.

In conclusion, our data show how human kinetochores can distinguish between changes in tension and microtubule occupancy. To achieve this the structure has in-built occupancy (Ndc80 jack-knife) and tension (Knl1 unravelling) sensors that would enable kinetochores to modulate their behaviour (*i.e*. spindle checkpoint and error correction processes) in response to changing mechanical inputs.

## Author Contributions

Project conception, planning and supervision was carried out by AM and NJB. All experiments were carried out by ER, except the Photoactivation experiments which were carried out by AT. Analysis of the intra-kinetochore measurement experiments was carried out by ER with help from TG, while ME generated Ndc80-EGFP knock-in cell lines. AM, NJB, ER and TG interpreted the data. CS developed kinetochore analysis code, NJB developed the Euclidean distance correction algorithm and PE built the visualizations of kinetochore organization. Manuscript preparation were carried out by AM and NJB with input from ER and TG.

## Acknowledgements

We thank Onur Sen for help analysing the MC191 cell line, CAMDU (Computing and Advanced Microscopy Unit) for support, Andrea Musacchio, Iain Cheeseman for antibodies, Jonathon Pines for cell lines and Jonathan Harrison for proof. This work was supported by the Leverhulme Trust (RPG-2017-349 to ADM and NJB). Work in the ADM lab was also supported by a Wellcome Trust Senior Investigator Award (106151/Z/14/Z) and a Royal Society Wolfson Research Merit Award (WM150020). TG and AT are funded by the Medical Research Council Doctoral Training Partnership grant MR/J003964/1.

---

**Supplemental Figure 1.**
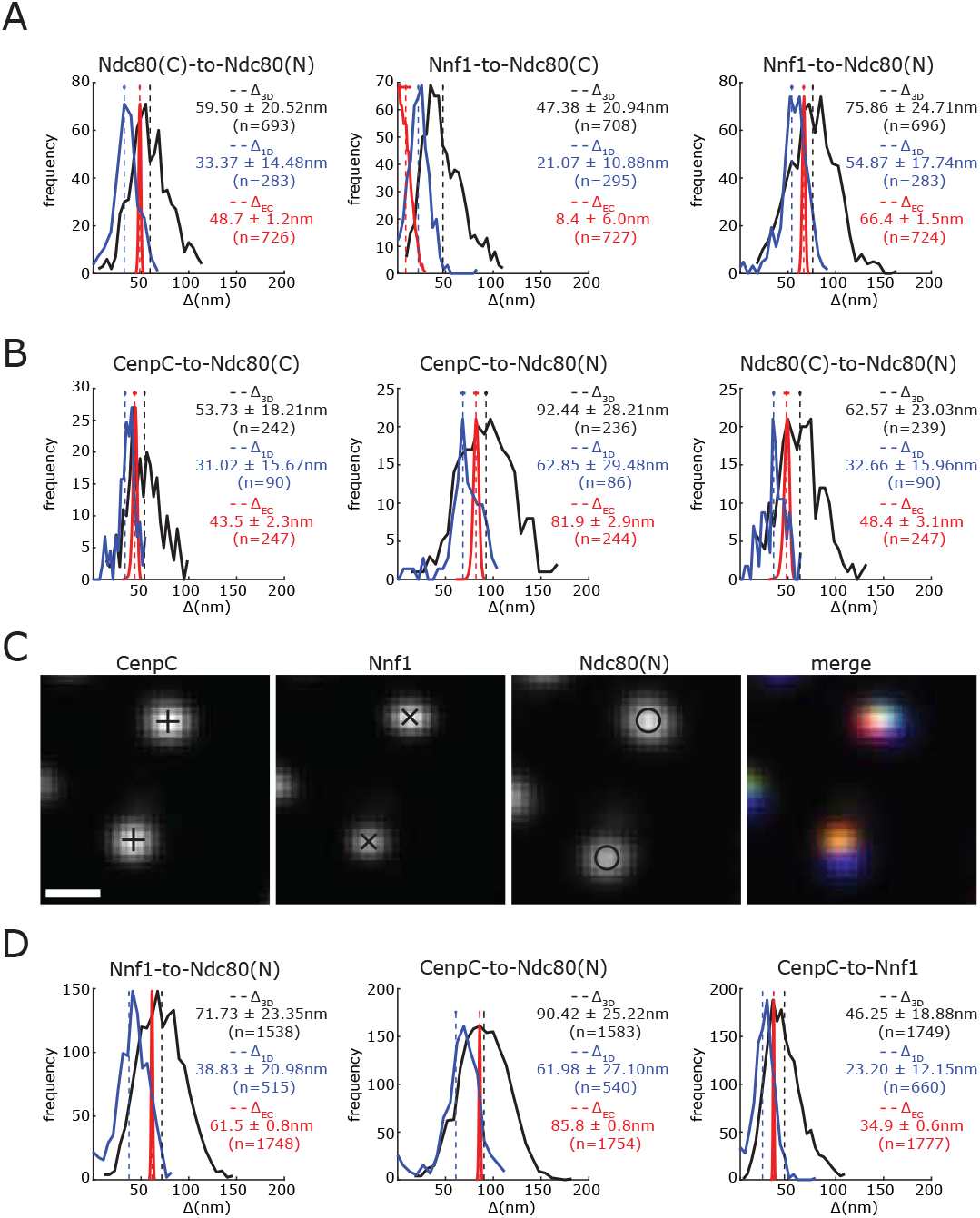
Comparison of Δ_1D_, Δ_3D_ and Δ_EC_ measurements. (A) Histograms showing the Ndc80(C)-to-Ndc80(N), Nnf1-to-Ndc80(C) and Nnf1-to-Ndc80(N) Δ_1D_, Δ_3D_ and the Δ_EC_ distances in RPE1 Ndc80-EGFP cells. (B) Histograms showing the CenpC-to-Ndc80(C), CenpC-to-Ndc80(N) and the Ndc80(C)-to-Ndc80(N) Δ_1D_, Δ_3D_ and the Δ_EC_ distances in RPE1 Ndc80-EGFP cells. (C) Kinetochore pair in RPE1 cells stained with anti-CenpC, anti-Nnf1 and anti-Hec1(9G3) antibodies. Scale bar 500 nm. (D) Histograms show the Nnf1-to-Ndc80(N), CenpC-to-Ndc80(N) and CenpC-to-Nnf1 Δ_1D_, Δ_3D_ and the Δ_EC_ distances in RPE1 cells. Histogram legends indicates distribution mean±sd. Vertical lines indicate means, bars on top dash show sd of estimated mean.

**Supplemental Figure 2.**
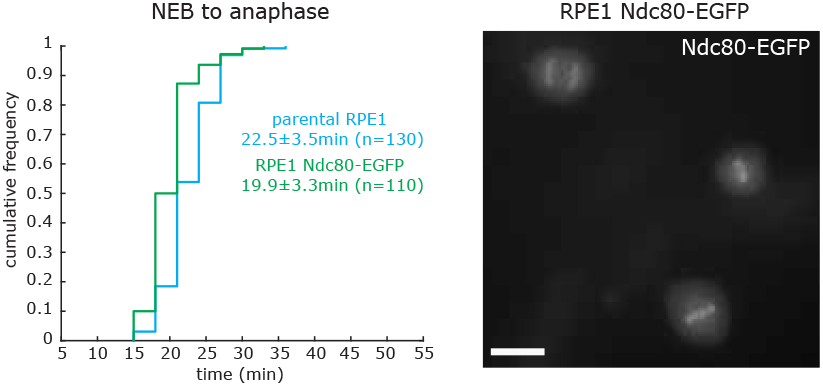
Parental and Ndc80-EGFP expressing RPE1 cells display the same mitotic timing. Cumulative frequency plot displaying the timing between nuclear envelope breakdown (NEB) to anaphase onset in parental RPE1 and RPE1 Ndc80-EGFP cells (MC191). Legend indicates sample mean±sd. Image on the right shows mitotic RPE1 cells expressing Ndc80-EGFP. Scale bars 20 *μ*m.

**Supplemental Figure 3.**
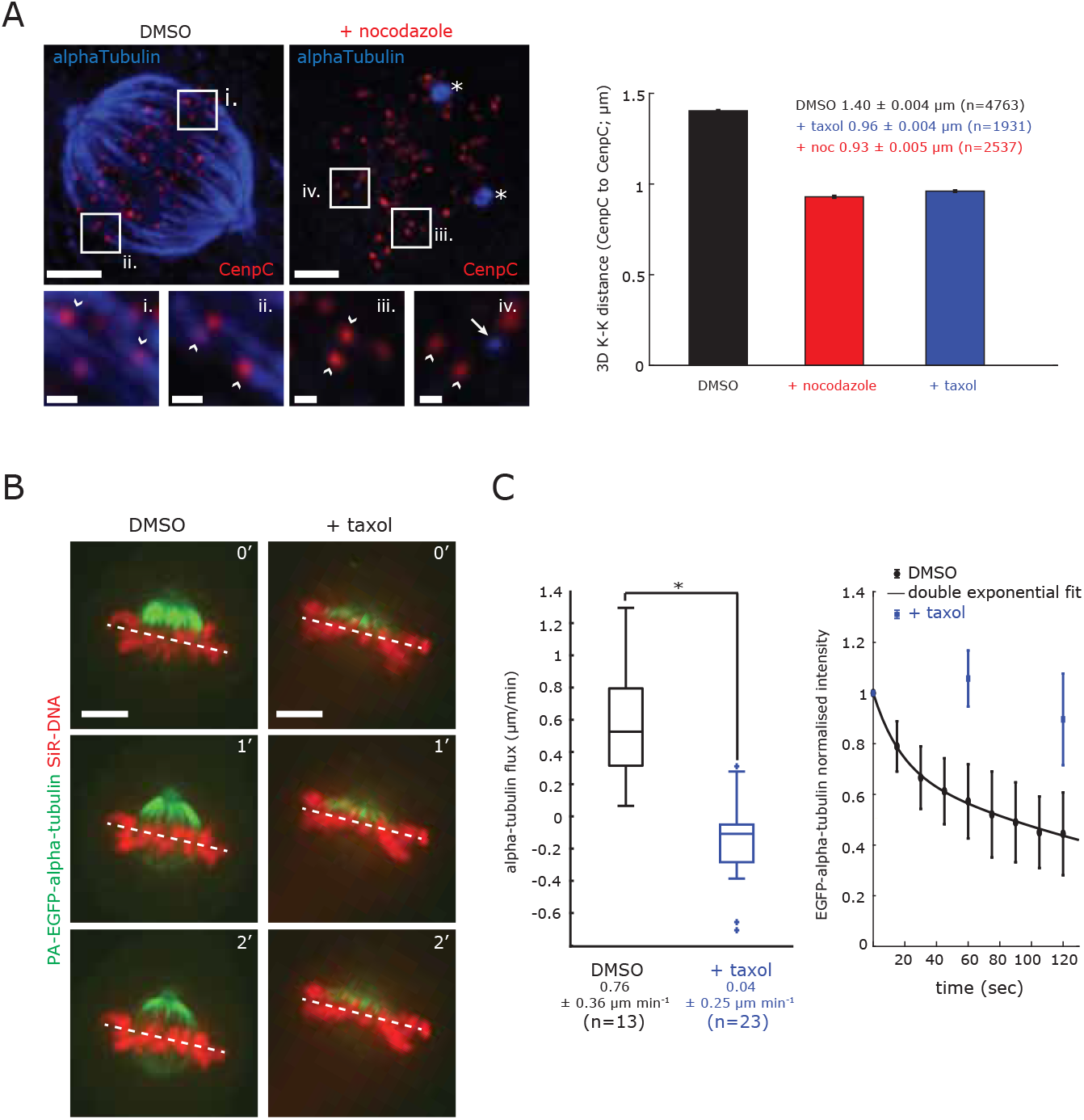
Nocodazole and taxol reduce inter-sister kinetochore distance and the taxol-dependent loss of inter-sister kinetochore tension is associated to reduced microtubule poleward flux. (A) Example images of RPE1 cells treated with 3.3 *μ*M nocodazole for 2 hr and DMSO as control. Scale bars 3 *μ*m. In control cells, bi-orientated kinetochore pairs (red) are attached to microtubules (blue) and are under tension (white arrowheads in insets i. and ii.; these images were enhanced to better show kinetochores and microtubules). Nocodazole induces full depolymerization of microtubules and reduces the inter-kinetochore distance (white arrowheads in inset iii. and iv.). In this condition, only centrosomes (*) and few microtubule stubs (white arrow in inset iv.) remain intact. Scale bar in insets is 500 nm. Bar charts (right) show the 3D K-K distance from CenpC to CenpC in cells treated with 3.3 *μ*M nocodazole for 2 hr, 1 *μ*M taxol for 15 min and DMSO as control. Legend indicates mean±SEM. (B) Example images RPE1 expressing photoactivatable-(PA)-EGFP-alpha-tubulin (green) stained with SiR-DNA to visualize the chromosomes (red). Photo-activation was carried out at T=0 and cells were imaged every 2 min. White dotted lines indicate the centre of the metaphase plate used as reference to measure the position of the PA-EGFP-alpha-tubulin (green). Scale bar 5 *μ*m. (C) Left panel: Box and whiskers plot showing the measured microtubule poleward flux in cells treated with 1 *μ*M taxol or DMSO for 15 min (see Materials and Methods for details). T-test indicates the difference is significant with 95% confidence interval: (*) p=3.5×10^−8^. Legend indicates sample mean±sd. Right panel: Intensity of PA-EGFP-alpha-tubulin at the indicated times in 1 *μ*M taxol or DMSO-treated cells (see Materials and Methods for details). Black solid line indicates the double exponential fitting for the DMSO intensities.

**Supplemental Figure 4.**
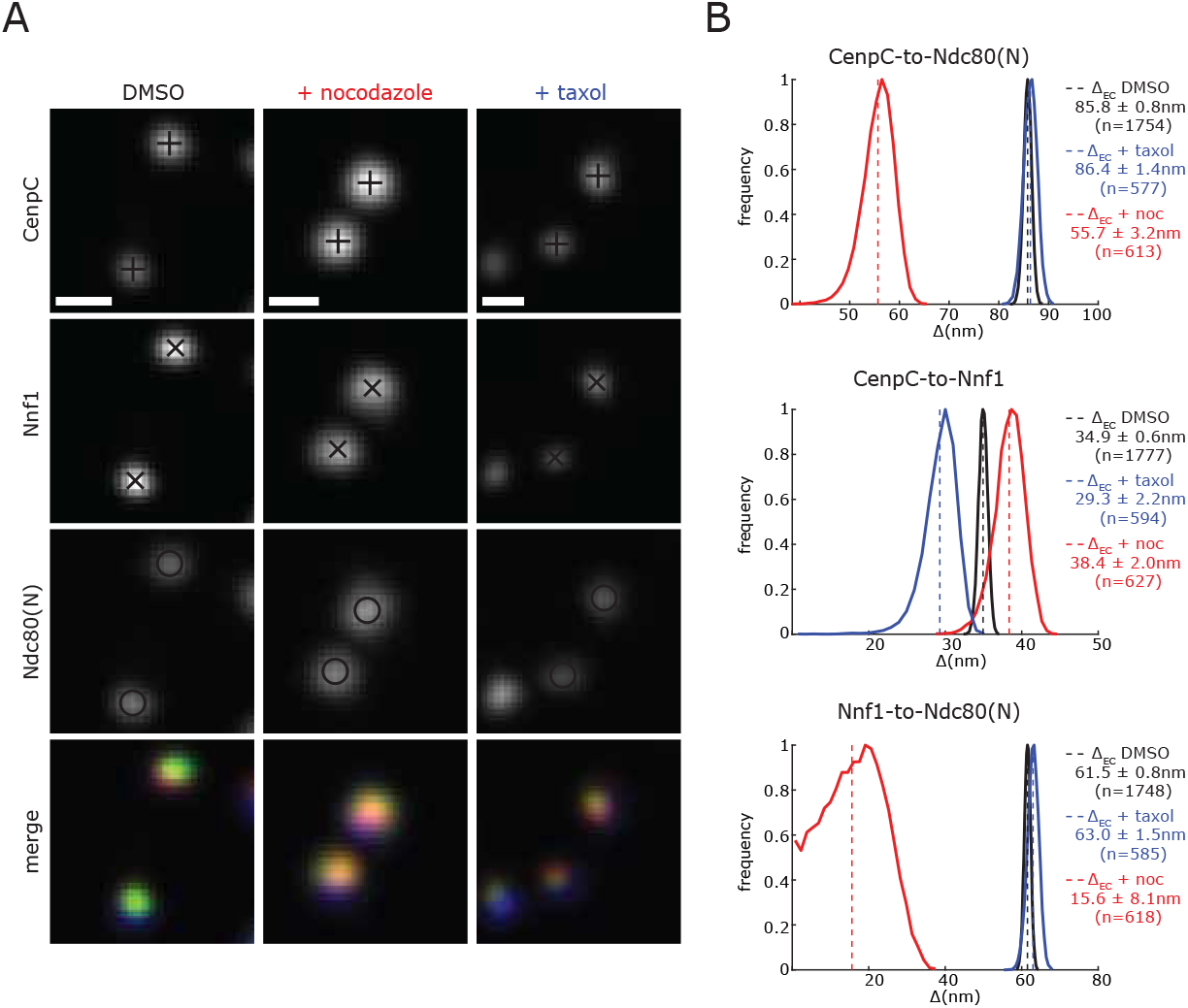
The Ndc80 N-terminus moves close to Nnf1 in unattached kinetochores. (A) Kinetochores in RPE1 cells stained with anti-CenpC, anti-Nnf1 and anti-Hec1(9G3) antibodies and treated with 3.3 *μ*M nocodazole for 2 hr, 1 *μ*M taxol for 15 min and DMSO as control. Scale bar 500 nm. (B) Distribution of the Nnf1-to-Ndc80(N), CenpC-to-Nnf1 and CenpC-to-Ndc80(N) Δ_EC_ distances in DMSO, nocodazole (+noc) and taxol treated cells. Legend indicates distribution mean±sd.

**Supplemental Figure 5.**
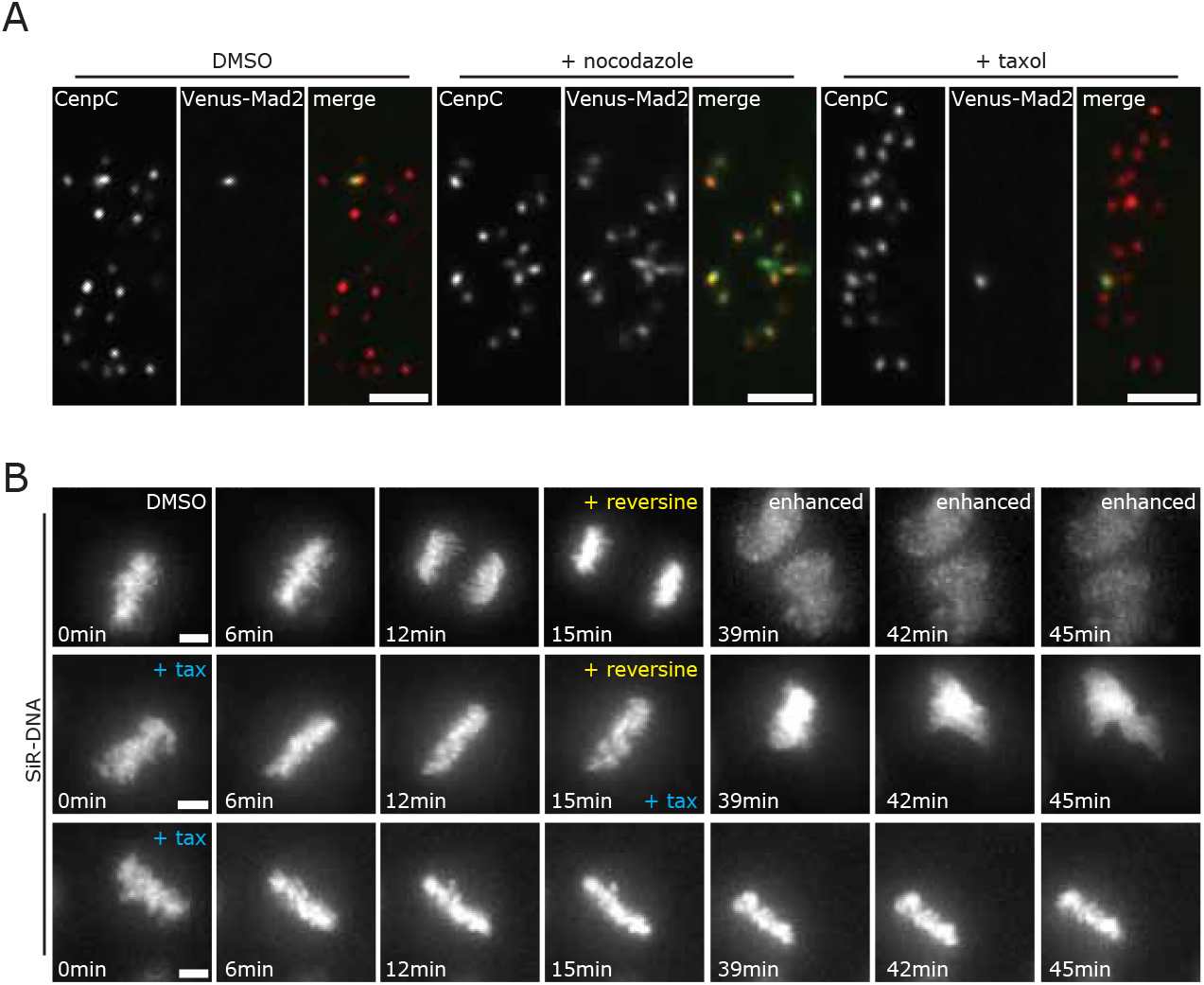
Taxol-induced mitotic delay is dependent on the activation of the Spindle Assembly Checkpoint. (A) Images of RPE1 Venus-Mad2 cells stained with anti-CenpC antibody and treated with 3.3 *μ*M nocodazole for 2 hr, 1 *μ*M taxol for 15 min and DMSO as control. Scale bars 3*μ*m. (B) Images of RPE1 cells treated with 1 *μ*M taxol (+tax) or DMSO for 15min and then with 1 *μ*M reversine for 2h. Bottom row shows a cell treated with 1 *μ*M taxol (+tax) only, as negative control. Time frame images at 39, 42 and 45 min in the DMSO sample were enhanced to better display the cell exiting mitosis. To visualize DNA, cells were previously treated with SiR-DNA. Scale bars 5 *μ*m.

**Supplemental Figure 6.**
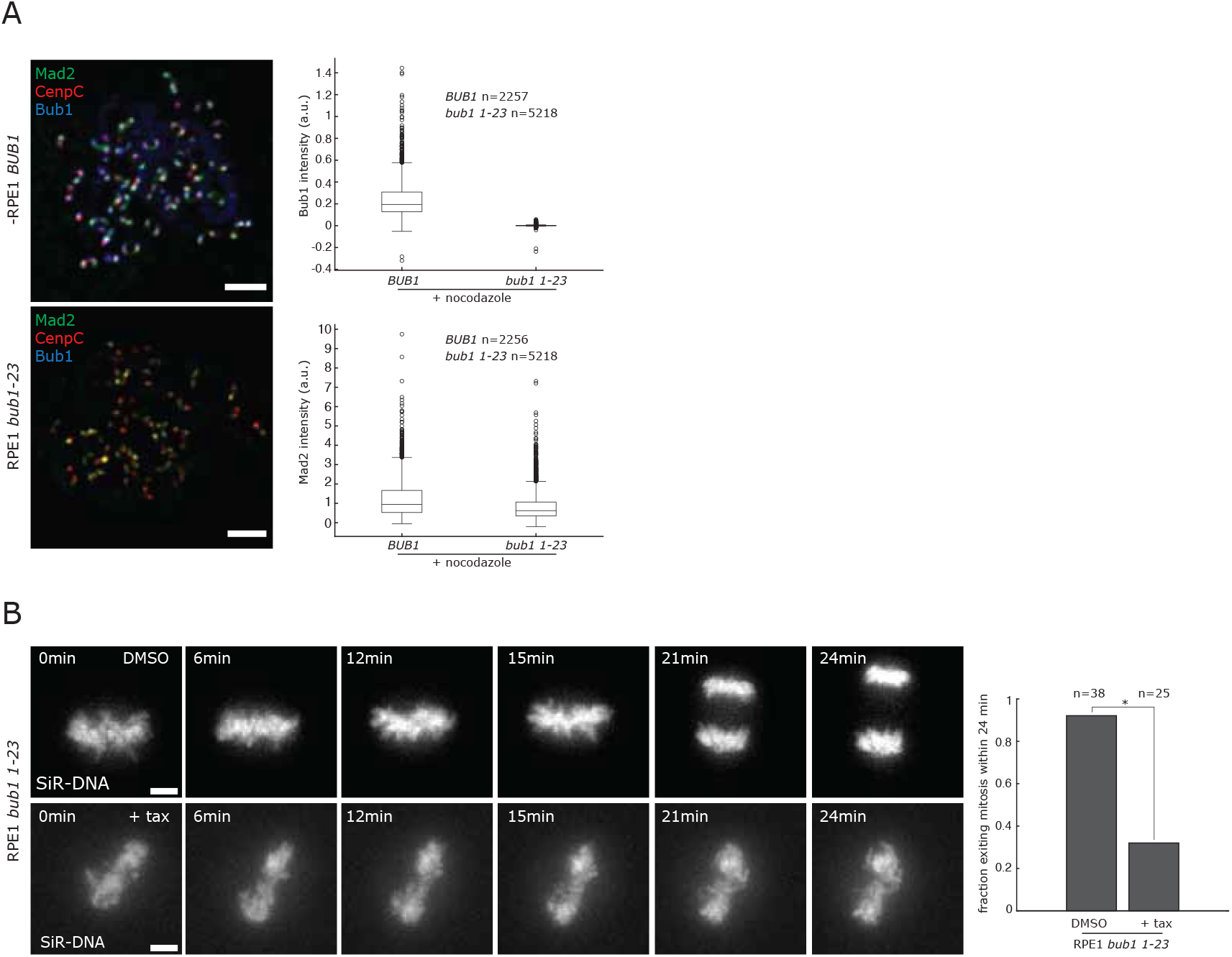
*bub1 1-23* cells can recruit Mad2 at kinetochores and are delayed in metaphase upon taxol treatment. (A) Images of parental and RPE1 *bub1 1-23* cells stained with anti-CenpC, anti-Bub1 and anti-Mad2 antibodies and treated with 3.3 *μ*M nocodazole for 2 hr. Scale bars 500 nm. Box and whiskers plots on the right show Bub1 and Mad2 intensities in parental and RPE1 *bub1 1-23* cells. Bub1 and Mad2 signals were background-subtracted and normalised to CenpC signal (also background corrected). (B) Images of RPE1 *bub1 1-23* cells treated with 1 *μ*M taxol (+tax) and DMSO as control, for 24 min. To visualize DNA, cells were previously treated with SiR-DNA. Scale bars 3 *μ*m. Bar chart on the right shows the fraction of cells exiting mitosis within 24 min. Fisher’s exact test indicates the differences are significant with 99% confidence interval: (*) p=7×10^−7^.

**Supplemental Table 1.**

| Distance |  | treatment | Δ1D mean ¥ | sd † | n | Δ3D mean | sd † | n | ΔEC mean | sd * | n | cell line |
| --- | --- | --- | --- | --- | --- | --- | --- | --- | --- | --- | --- | --- |
| CenpC-A488 | CenpC-A568 | untreated | -0.62 | 9.28 | 1111 | 31.60 | 17.64 | 2279 | 2.2 | 1.5 | 2302 | RPE1 |
| CenpC-A488 | CenpC-A647 |  | -3.33 | 12.77 | 631 | 45.61 | 23.29 | 1364 | 4.3 | 2.7 | 1437 |  |
| CenpC-A568 | CenpC-A647 |  | -2.74 | 10.98 | 653 | 43.39 | 21.77 | 1388 | 4.5 | 2.8 | 1452 |  |
| Ndc80(C) | Ndc80(N) | DMSO | 33.37 | 14.48 | 283 | 59.50 | 20.52 | 693 | 48.7 | 1.2 | 726 | RPE1 Ndc80-EGFP (MC191) |
|  |  | nocodazole | 1.12 | 19.04 | 227 | 50.57 | 23.10 | 669 | 7.8 | 5.0 | 694 |  |
|  |  | taxol | 27.68 | 12.86 | 217 | 59.96 | 21.42 | 582 | 45.5 | 1.4 | 620 |  |
| Nnf1 | Ndc80(N) | DMSO | 54.87 | 17.74 | 283 | 75.86 | 24.71 | 696 | 66.4 | 1.5 | 724 |  |
|  |  | nocodazole | 16.04 | 19.74 | 218 | 56.14 | 24.82 | 657 | 12.7 | 7.2 | 696 |  |
|  |  | taxol | 49.17 | 15.01 | 218 | 74.42 | 21.91 | 587 | 68.3 | 1.2 | 624 |  |
| Nnf1 | Ndc80(C) | DMSO | 21.07 | 10.88 | 295 | 47.38 | 20.94 | 708 | 8.4 | 6.0 | 727 |  |
|  |  | nocodazole | 14.64 | 16.47 | 229 | 51.41 | 22.62 | 671 | 8.7 | 5.6 | 699 |  |
|  |  | taxol | 22.76 | 11.81 | 226 | 52.31 | 22.93 | 596 | 11.3 | 7.0 | 627 |  |
| Ndc80(C) | Ndc80(N) | DMSO | 32.66 | 15.96 | 90 | 62.57 | 23.03 | 239 | 48.4 | 3.1 | 247 | RPE1 Ndc80-EGFP (MC191) |
|  |  | nocodazole | 6.09 | 16.02 | 149 | 52.15 | 41.32 | 561 | 7.3 | 4.6 | 600 |  |
|  |  | taxol | 25.09 | 14.25 | 238 | 59.91 | 21.56 | 677 | 44.4 | 1.6 | 731 |  |
| CenpC | Ndc80(N) | DMSO | 62.85 | 29.48 | 86 | 92.44 | 28.21 | 236 | 81.9 | 2.9 | 244 |  |
|  |  | nocodazole | 42.19 | 20.37 | 150 | 69.87 | 23.99 | 544 | 56.6 | 2.0 | 597 |  |
|  |  | taxol | 58.49 | 17.91 | 245 | 87.57 | 22.93 | 683 | 83.0 | 1.1 | 732 |  |
| CenpC | Ndc80(C) | DMSO | 31.02 | 15.67 | 90 | 53.73 | 18.21 | 242 | 43.5 | 2.3 | 247 |  |
|  |  | nocodazole | 36.79 | 16.48 | 154 | 67.53 | 38.31 | 563 | 40.4 | 9.3 | 599 |  |
|  |  | taxol | 33.13 | 12.80 | 254 | 58.55 | 20.09 | 695 | 48.9 | 1.1 | 734 |  |
| Nnf1 | Ndc80(N) | DMSO | 38.83 | 20.98 | 515 | 71.73 | 23.35 | 1538 | 61.5 | 0.8 | 1748 | RPE1 |
|  |  | nocodazole | 12.54 | 17.59 | 126 | 62.76 | 26.03 | 513 | 15.6 | 8.1 | 618 |  |
|  |  | taxol | 37.48 | 17.92 | 152 | 74.55 | 14.13 | 491 | 63.0 | 1.5 | 585 |  |
| CenpC | Ndc80(N) | DMSO | 61.98 | 27.10 | 540 | 90.42 | 25.22 | 1583 | 85.8 | 0.8 | 1754 |  |
|  |  | nocodazole | 40.25 | 22.05 | 126 | 76.07 | 28.36 | 518 | 55.7 | 3.2 | 613 |  |
|  |  | taxol | 59.22 | 20.89 | 136 | 91.06 | 24.35 | 476 | 86.4 | 1.4 | 577 |  |
| CenpC | Nnf1 | DMSO | 23.20 | 12.15 | 660 | 46.25 | 18.88 | 1749 | 34.9 | 0.6 | 1777 |  |
|  |  | nocodazole | 29.00 | 15.39 | 185 | 55.14 | 22.66 | 603 | 38.4 | 2.0 | 627 |  |
|  |  | taxol | 21.21 | 10.86 | 195 | 46.37 | 19.85 | 576 | 29.3 | 2.2 | 594 |  |
| CenpC | Bub1(N) | DMSO | 37.81 | 21.07 | 902 | 72.03 | 25.41 | 2053 | 58.9 | 1.1 | 2232 | RPE1 |
|  |  | nocodazole | 44.27 | 22.69 | 360 | 70.39 | 23.38 | 1048 | 60.7 | 1.1 | 1137 |  |
|  |  | taxol | 36.53 | 25.84 | 190 | 82.45 | 44.60 | 684 | 63.1 | 2.8 | 816 |  |
| CenpC | Rod(C) | DMSO | 47.95 | 23.39 | 389 | 85.62 | 31.24 | 1073 | 71.4 | 1.9 | 1294 | RPE1 |
|  |  | nocodazole | 46.55 | 26.34 | 16 | 89.03 | 48.42 | 91 | 65.6 | 8.3 | 112 |  |
|  |  | taxol | 37.77 | 27.91 | 56 | 84.14 | 31.31 | 282 | 70.5 | 3.6 | 350 |  |
| CenpC | Zwlich | DMSO | 73.41 | 19.35 | 408 | 100.01 | 28.30 | 1090 | 96.3 | 1.0 | 1292 | RPE1 |
|  |  | nocodazole | 63.29 | 30.83 | 66 | 103.91 | 43.57 | 331 | 89.4 | 3.2 | 397 |  |
|  |  | taxol | 67.89 | 22.22 | 100 | 97.53 | 26.30 | 440 | 96.6 | 1.4 | 565 |  |
| CenpC | pKnl1(S24) | DMSO | 33.48 | 40.20 | 263 | 89.48 | 45.03 | 959 | 46.7 | 10.5 | 1198 | RPE1 |
|  |  | nocodazole | 68.87 | 43.85 | 47 | 115.75 | 52.84 | 346 | 94.8 | 3.6 | 421 |  |
|  |  | taxol | 86.12 | 39.89 | 51 | 124.92 | 43.96 | 409 | 115.2 | 2.3 | 492 |  |
| CenpC | Knl1 - MELT 2 | DMSO | 51.89 | 13.19 | 445 | 72.76 | 23.49 | 1012 | 65.4 | 1.0 | 1066 | RPE1 |
|  |  | nocodazole | 56.72 | 26.23 | 136 | 90.87 | 29.82 | 487 | 82.5 | 2.2 | 605 |  |
|  |  | taxol | 51.47 | 15.61 | 166 | 78.10 | 37.45 | 411 | 68.5 | 1.9 | 443 |  |
| CenpC | Mad2 | nocodazole *BUB1* | 51.97 | 19.68 | 105 | 80.07 | 26.25 | 478 | 70.9 | 1.8 | 550 | RPE1 |
|  |  | nocodazole *bub1 1-23* | 58.70 | 32.28 | 220 | 97.18 | 36.74 | 1016 | 84.1 | 1.8 | 1265 | RPE1 *bub1 1-23* (MC170) |
| CenpC | Ndc80(N) | DMSO - Mad2+ | 45.52 | 25.71 | 5 | 71.85 | 22.58 | 68 | 66.2 | 3.7 | 74 | RPE1 Venus-Mad2 |
|  |  | DMSO - Mad2- | 66.08 | 15.20 | 278 | 86.59 | 22.06 | 685 | 82.4 | 1.0 | 721 |  |
|  |  | DMSO - pool Mad2+/- | 63.76 | 16.70 | 334 | 85.26 | 22.50 | 753 | 81.0 | 1.0 | 795 |  |
|  |  | nocodazole - Mad2+ | 43.73 | 18.86 | 510 | 70.89 | 25.73 | 1218 | 58.3 | 1.3 | 1322 |  |
|  |  | taxol - Mad2+ | 45.06 | 39.48 | 5 | 84.33 | 26.96 | 54 | 80.5 | 4.9 | 64 |  |
|  |  | taxol - Mad2- | 69.63 | 14.14 | 194 | 94.78 | 23.92 | 498 | 92.0 | 1.3 | 558 |  |
|  |  | taxol - pool Mad2+/- | 68.42 | 15.44 | 234 | 93.76 | 24.41 | 552 | 90.9 | 1.3 | 622 |  |

¥ 1D measurements determined as Smith et al., 2016.
† sd of samples for data
\* sd of posterior (parameter sd)

**Supplemental Table 2.**

**Roscioli et al., Supplemental Table 2**
| Distance (vector) | | treatment | $\Delta$ expected (nm) † | ΔEC observed (nm) | nematic order (*N*) | markers labelled | cell line | |
| --- | --- | --- | --- | --- | --- | --- | --- | --- |
| Ndc80(C) | Ndc80(N) | DMSO | 50.08 | 48.7 | 0.97 | Nnf1-Ndc80(C)-Ndc80(N) | RPE1 Ndc80-EGFP (MC191) |  |
|  |  | nocodazole | 19 | 7.8 | 0.41 |  |  |  |
|  |  | taxol | 50.08 | 45.5 | 0.91 |  |  |  |
|  |  | DMSO | 50.08 | 48.4 | 0.97 | CenpC-Ndc80(C)-Ndc80(N) | RPE1 Ndc80-EGFP (MC191) |  |
|  |  | nocodazole | 19 | 7.3 | 0.38 |  |  |  |
|  |  | taxol | 50.08 | 44.4 | 0.89 |  |  |  |
| Nnf1* | Ndc80(N) | DMSO | 64.99 | 66.4 | 1.02 | Nnf1-Ndc80(C)-Ndc80(N) | RPE1 Ndc80-EGFP (MC191) |  |
|  |  | nocodazole | 34 | 12.7 | 0.37 |  |  |  |
|  |  | taxol | 64.99 | 68.3 | 1.05 |  |  |  |
|  |  | DMSO | 64.99 | 61.5 | 0.95 | CenpC-Nnf1-Ndc80(N) | RPE1 |  |
|  |  | nocodazole | 34 | 15.6 | 0.46 |  |  |  |
|  |  | taxol | 64.99 | 63.0 | 0.97 |  |  |  |
| Rod(C)‡ | Zwilch** | DMSO | 31.4 | 24.9±2.1 | 0.79±0.07 | CenpC-Rod(C) and CenpC-Zwilch | RPE1 |  |
|  |  | nocodazole | 31.4 | 23.8±8.9 | 0.76 ±0.28 |  |  |  |
|  |  | taxol | 31.4 | 26.1±3.9 | 0.83±0.12 |  |  |  |

\* distances estimated considering the Nnf1 antibody binding the central region of the MIS12 complex
\*\* distances estimated considering the Zwilch antibody binding the central region of the protein
† vector calculated considering Ndc80 hinge angle of 203.5° for DMSO and taxol and 0° for nocodazole samples
‡ distances between Zwilch and Rod derived by subtracting the CenpC-to-Rod(C) from CenpC-to-Zwilch distances

## Materials and Methods

### Cell culture

Immortalized (hTERT) diploid human retinal pigment epithelial (RPE1) cells, RPE1 *bub1 1-23* (MC170; (Currie et al., 2018)), RPE1 Ndc80-EGFP (MC191), RPE1 Venus-Mad2 (Mad2L1 Venus/+; kind gift from Jonathan Pines) and RPE1 expressing photoactivatable-(PA)GFP-alpha-tubulin (MC021, Toso et al., 2009) were grown in DMEM/F-12 medium containing 10% fetal bovine serum (FBS), 2 mM L-glutamine, 100 U/ml penicillin and 100 μg/ml streptomycin. 200 μg/ml Geneticin (G418, Invitrogen) was added to the media to maintain MC021 cells. All cell cultures were maintained at 37°C with 5% CO_2_ in a humidified incubator.

### CrispR-Cas9 genome editing

Small guide RNAs (sgRNAs) (5’-caccgATGCATGTCAGAAGATCTCT-3’ and 5’-aaacAGAGATCTTCTGACATGCATc-3’) targeting exon 17 of the *NDC80* gene were designed using http://crispr.mit.edu to insert a EGFP in frame after the stop codon. sgRNAs were annealed and ligated into pX330 which enables their expression in mammalian cells along with a humanized *S. pyogenes* Cas9 (Addgene). A homology Directed Repair (HDR) construct was designed with 800bp homology upstream of the Stop codon and 800 bp downstream of the stop codon. 1 μg of sgRNA construct and 1.5 μg of linearized HDR plasmid were transfected into RPE1 cells using Fugene 6 (Promega). Positive cells were FACS sorted to isolate the Ndc80-EGFP expressing cells and single clones were identified by visual inspection with fluorescent microscope, with a further round of clonal selection used to eliminate heterogeneity in the population. PCR analysis of clone MC191 confirmed the presence of a wildtype and EGFP-containing allele (primers: Fwd_Ndc80EGFP, 5’-TAAACTGCAGCCATATGTAGTAAC-3’; Rev_Ndc80EGFP, 5’-TTGAAATTAGTAAGAAATGAGAGA-3’). Individual alleles were then analysed by cloning PCR products and sequencing the products (primers: EGFPNtRev 5’-CCGGACACGCTGAACTTG-3’ for the EGFP containing allele band and Rev_Ndc80-EGFP 5’-TTGAAATTAGTAAGAAATGAGAGA-3’ for the wild type allele band). This confirmed that the EGFP was in-frame with the 3’-end of *NDC80*, although a single amino acid change (Threonine 635 to Alanine, T635A) was identified in the unstructured tail distal to the coiled coils that are required for NDC80 complex tetramerisation. We also note that this varient is found in all primates, except *H. sapiens* and *H. Neanderthalensis*, and that no difference in mitotic timing or multiple delta measurements (Supplementary Table 1) were detected when compared to parental RPE1 cells (Supplemental Figure 2).

### Fixed cell experiments and immunofluorescence

Cells were plated on glass coverslips (0.16-0.19 mm) 24 or 48 hr before treatment with 3.3 *μ*M nocodazole (Sigma, diluted in DMSO) for 2 hr, 1 *μ*M taxol (Sigma, diluted in DMSO) for 15 min or with 0.001% DMSO for 2 hr as a control (Sigma). Cells were then fixed in 10 mM EGTA (Sigma), 1 mM MgCl_2_ (Sigma), 20 mM PIPES pH 6.8 (Sigma), 0.2% Triton-X100 (Fisher), and 4% formaldehyde (VWR Life Science) for 10 min, washed 3 times in PBS before incubation in PBS supplemented with 3% BSA (Sigma) for 30 min to block non-specific antibody binding. Next, cells were incubated with primary antibodies for 1 hr, washed 3 times in PBS and then incubated for 30 min with secondary antibodies and DAPI (Sigma, 1:1000 dilution); all antibodies were diluted in PBS + 3% BSA and are as follows: Primary antibodies - guinea pig anti-CenpC (1:2000, MBL, PD030), mouse anti-Hec1(9G3) (1:1000, abcam, ab3613), mouse anti-Bub1 (1:200, abcam, ab54893), mouse anti-Rod (1:50, abcam, ab56745), mouse anti-alpha tubulin (1:1000, Sigma, T6074), rabbit anti-Zwilch (1:1000, gift from Andrea Musacchio), rabbit anti-Nnf1 (1:1000, McAinsh et al, 2006), rabbit anti-pKnl1(S24) (1:2200, gift from Iain Cheeseman), rabbit anti-Knl1 (1:500, abcam, ab70537), rabbit anti-Mad2 (1:500, BioLegend, Poly19246). Secondary antibodies - AlexaFluor-488nm goat anti-mouse, got anti-rabbit and goat anti-guinea pig (1:500; Invitrogen), AlexaFluor-568nm goat anti-guinea pig (1:500; Invitrogen), AlexaFluor-594nm goat anti-rabbit (1:500; Invitrogen) and AlexaFluor-647nm goat anti-mouse and goat anti-guinea pig (1:500; Invitrogen). Cells were then washed in PBS and mounted in Vectashield (Vector).

### Fixed cells image acquisition and processing

Image stacks were acquired using a confocal spinning-disk microscope (VOX UltraView; PerkinElmer, UK) equipped with a 100X / 1.4 NA oil-immersion objective and a Hamamatsu ORCA-R2 camera, controlled by Volocity 6.0 (PerkinElmer) running on a Windows 7 64-bit (Microsoft, Redmond, WA) PC (IBM, New Castle, NY). Image stacks were acquired over 61 z-slices separated by 0.2 *μ*m using the 488, 561, 640 and 405 nm wavelength lasers. Acquisition settings were set in order that the kinetochore signals were 100-150 units above background. Spinning disc images were exported from Volocity 6.0 in OME.TIFF format (The Open Microscopy Environment, UK) and deconvolved using Huygens 4.1 (SVI), using point spread functions (PSFs) calculated from 100 nm TetraSpeck fluorescent microspheres (Invitrogen) using the Huygens 4.1 PSF distiller. Where required images in the 640 nm wavelength were deconvolved within Huygens 4.1 using a theoretical PSF. Deconvolved images were exported from Huygens 4.1 in r3d format (Applied Precision, Slovakia) and read into MATLAB (2017a, 2018a, Mathworks, Natick, MA) using the loci-tools java library (The Open Microscopy Environment). Kinetochores spots were first detected using the 561 nm channel and then (where appropriate) signals from secondary and tertiary markers in the 488 and 640 nm channels identified (Smith et al., 2016). Correction of chromatic aberrations for all three channels was carried out using images taken from a reference slide (either RPE1 HaloTag-CenpA labelled with Oregon Green, TMR and JF646; or anti-CenpC staining with a mix of alexa488, 568 and 647-labelled secondary antibodies) on the same day as acquisition for experiments (Dudka et al., 2018; Smith et al., 2016). All kinetochore tracking, sister pairing, 3D intra-kinetochore distances and intensity measurements were made using KiT (<u>K</u>inetochore <u>T</u>racking) v2.1.10 with additional functions (see: https://github.com/cmcb-warwick; (Olziersky et al., 2018)). Bub1, Mad2 and pKnl1 (S24) signal intensities were background subtracted and then normalised using the CenpC signal (also background subtracted), whereas for measurement of endogenous Venus-Mad2 the non-normalised signal is reported.

### Analysis of Spindle Assembly Checkpoint activation

Cells were cultured in four compartment CELLview dish (627975, Greiner Bio-One Ltd.). Time-lapse imaging was performed on an Olympus DeltaVision microscope (Applied Precision, LLC) equipped with a Photometrics CoolSNAP HQ camera (Roper Scientific) and a stage-top incubator (TokaiHit) to maintain cells at 37°C and 5% CO_2_. Temperature was further stabilized using a microscope enclosure (Weather station; Precision Control) held at 37°C. Image stacks (7 × 2 *μ*m optical sections) were acquired using the softWoRX 6.0 software every 3 min using a 40x / 1.3 NA oil-immersion objective. To visualise the DNA, cells were incubated, 1 hr before imaging, with DMEM/F-12 media containing 0.5 *μ*M SiR-DNA (Spirochrome). In each experiment, only fields (1024 × 1024 pixels) containing at least one metaphase cells were imaged using the point visit function in softWoRX 6.0. Imaging started after the addition of DMSO or 1 *μ*M taxol-containing media. Cells were imaged for 15 min to reproduce the same conditions used for the fixed cell experiments. For experiments with RPE1 *bub1 1-23* cells imaging was extended to 24 min because the timing to anaphase onset was slightly delayed with respect to the parental cell line. In the experiments with reversine, cells were treated with either DMSO or 1 *μ*M taxol-containing media for 15 min and then 1 *μ*M reversine (Sigma)-containing media was added for 120 min (total imaging time was 135 min). As control, cells were treated with 1 *μ*M taxol and imaged for 135 min. Images were acquired at 32 % neutral density using Cy5 filter and an exposure time of 0.05 s. Timing of exit from mitosis was scored by eye.

### Mitotic timing analysis

Parental RPE1 and RPE1 stably expressing Ndc80-EGFP were cultured in glass bottom FluoroDish (FD35-100, World Precision Instrument, Inc.). Time-lapse imaging was performed on Olympus DeltaVision microscopes (Applied Precision, LLC) equipped either with Photometrics CoolSNAP HQ (Roper Scientific) or Photometrics CoolSNAP HQ2 cameras (Roper Scientific) and temperature held at 37°C as described above. Image stacks (7 × 2 *μ*m optical sections) were acquired using the SoftWoRX 6.0 software every 3 min using a 40x / 1.3 NA oil-immersion objective. To visualise the DNA, RPE1 cells were incubated, 1 hr before imaging, with DMEM/F-12 media containing 0.5 *μ*M SiR-DNA (Spirochrome). In each experiment, 30 to 40 fields (1024 × 1024 pixels) were imaged using the point visit function in softWoRX 6.0. Images were acquired for 15 hr at 10% neutral density using Cy5 filter and an exposure time of 0.05 s. The timing of nuclear envelope breakdown and anaphase onset were scored by eye.

### Microtubule poleward flux and turnover analysis

RPE1 cells stably expressing photoactivatable-(PA)GFP-alpha-tubulin (Toso et al., 2009) were cultured in Fluorodishes (FD35-100, World Precision Instrument, Inc.) and DNA visualised by incubation for 30 min, with CO_2_ independent L15 media (Invitrogen, UK) containing 0.5 *μ*M SiR-DNA (Spirochrome, CH). DMSO or 1*μ*M taxol was added 15 min prior to imaging. Photoactivation was carried out using a confocal spinning disk microscope (Marianas SDC, 3i, UK) equipped with a Vector module for photoactivation, a 100x / 1.46 NA immersion oil objective and a Photometrics 95B Prime sCMOS camera controlled by Slidebook 6.0 (3i, UK). Cells were maintained at 37°C using a stage top incubator (both Okolab, Italy). PAGFP-tubulin was activated in an ROI (100×5 pixels, parallel to the metaphase plate) with 4 × 2 ms pulses of a 405 nm laser. Images were then acquired (excitation 488 nm) every 15 s for 2 min (150 ms exposure, 3 planes 1 *μ*m z-step centered on the photoactivated plane). Images of the DNA staining were acquired (excitation 640nm) at frame 1, 5 and 9. Poleward flux was calculated by measuring the displacement of photoactivated marks over the first 5 frames. To determine the turnover of PAGFP-alpha-Tubulin the background-subtracted pixel intensity of the photo-activated EGFP-alpha-tubulin over time was measured (averaging the mean intensity of two 7×7 pixels square boxes placed on the photo-activated region). In DMSO, intensity was measured at every time point, whereas, in taxol, intensity was analysed at time points 0 min, 1 min and 2 min. Intensity measurements were done by.

## Supplemental Methods

Supplemental Methods contains details of 1) the Bayesian Euclidean distance correction algorithm, 2) Kinetochore architecture simulations and 3) Nematic order.

## Supplemental Methods: Roscioli et al

### 1 A Bayesian Euclidean distance correction algorithm

#### 1.1 Introduction

From multiple samples of intra-kinetochore distance measurements between two fluorophores we present a Bayesian algorithm to determine the 3D distance. Noise in the measurement means that the measured distance is an over-estimate, e.g. see (Churchman et al., 2005, 2006), a bias that needs to be corrected. Measurement noise comes from the microscope point spread function (PSF) and the brighness of the flourophores, so is both wavelength dependent and protein density (flourophore density and brightness) dependent. Isotropic measurement errors have been analysed before with available formulae/maximum likelihood correction algorithms in 1-3D (Churchman et al., 2006). However, non-isotropic measurement errors have not been analysed.

In section 1.2 we introduce a 3D model of paired fluorophore measurements, in section 1.3 we derive the associated likelihood. In section 1.4 we give the updates for our MCMC sampler. In section 1.5 we demonstrate performance on simulated data.

#### 1.2 Euclidean distance model

Let 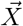 be the observed vector from flourophore 1 to fluorophore 2. For *N* measurements, we have samples 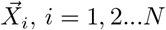. Assume that the measurement noise is Gaussian, and if the true displacement is 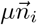, direction 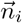 (unit vector) and true distance *μ*, we have

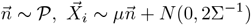

where 0.5Σ is the 3D precision matrix (inverse of covariance) for the spot centre accuracy, and direction vector 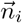 is chosen according to a direction distribution 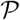, e.g. uniform over a sphere. The task is to determine the mean distance *μ* ≥ 0 and the measurement error covariance matrix 2Σ^−1^. The factor of 2 comes from the fact that both fluorophores have measurement error, so we could write this as the sum of the two measurement errors 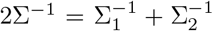; however we don’t estimate these separately. For isotropic measurements Σ is diagonal; 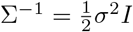, *I* is the diagonal matrix and σ is the intradistance error standard deviation. Thus, if spot accuracies are identical, individual spot centres would have variance *σ*^2^/2. Note that the measurement noise can reverse the orientation, i.e. 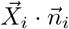 can be negative.

To quantify distances, it is natural to consider the distribution of 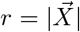; this involves integrating over 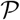 and the angular components of 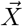 (i.e. angular measurements are ignored). For the case of isotropic errors we take advantage of the rotational invariance, i.e. choose axes relative to 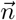; thus, the only coordinate is *r* - the distance between the fluorophores. Then, for isotropic measurement error in 3D we obtain the probability density (Churchman et al., 2006),

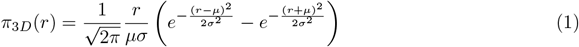

This gives for the mean and variance,

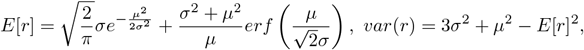

which can be fitted to data and μ,σ thereby inferred

#### 1.3 Likelihood based inference

For the non-isotropic measurement error case there is no analytical form as above, as far as we are aware. Thus, we use a simulation methodology to sample from the posterior distribution that is based on the likelihood. For *N* 3D measurements the likelihood is given by,

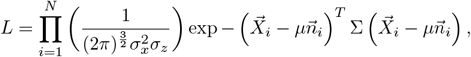

where 2^−1^Σ is the 3D precision matrix for the spot centre accuracy, and 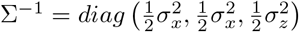 in microscope coordinates.

We use spherical coordinates to specify the relative position of the two vectors 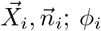 is the relative angle in the *x, y* plane between 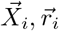, and *θ_i_, θ_Xi_* are the angles of 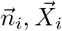 with the *z*-axis:

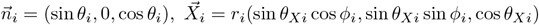

Then the likelihood reads,

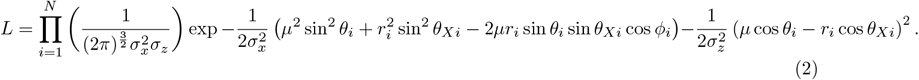

This has parameters *μ,σ_x_,σ_z_* and sample specific hidden (unmeasured) variables *θ_i_,ϕ_i_, i* = 1, 2…*N* to determine (recall *r_i_, θ_Xi_* are measured). Then using the spherically symmetric measure (solid angle) *d*Ω = sin *θdθdϕ* and Bayes theorem, the posterior distribution is given by,

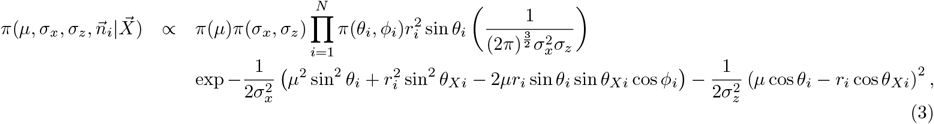

where *π*(*μ*), *π*(*σ_x_,σ_z_*) and *π*(*θ_i_,ϕ_i_*) are appropriate priors. If 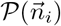 is uniform over the surface of the sphere the prior on cos *θ_i_* is *U*([1, –1]).

This distribution has 3 global variables *μ, σ_x, z_*, and 2*N* hidden variables *ϕ_i_,θ_i_*. To generate samples from this posterior we use a Markov chain Monte Carlo (MCMC) algorithm. However, the posterior can be marginalised in *ϕ_i_* (integrating out *ϕ_i_* using a uniform prior) to give the alternative form with only *N* hidden variables *θ_i_*,

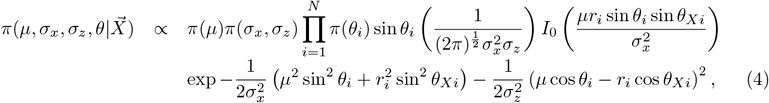

where *I*_0_ is the modified Bessel function. This form is computationally more expensive per step but has superior mixing so convergence is achieved with fewer steps. This is the form we use.

If the measurement error is isotropic we can also integrate out the angular variables *θ_i_* to obtain the posterior corresponding to (1).

#### 1.4 MCMC sampling algorithm

We use an algorithm that updates variables separately except for *μ, σ_x_* that are highly correlated. We switch to precisions 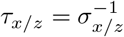 as is typical for models with Gaussian noise since precisions are (conditionally) Gamma distributed under choice of a conjugate Gamma prior. We use short hand *s_i_* = sin*θ_i_*, *c_i_* = cos*θ_i_*, *s_Xi_* = sin*θ_Xi_*, *s_Xi_* = sin*θ_Xi_*. We use weak conjugate priors, imposing any positivity conditions by truncation. Updates are as follows:

##### Joint *μ, τ_x_* proposal

We find that *μ* and *τ_x_* are often highly correlated. Here we describe a twisted random walk along the eigendirections of the covariance matrix. For estimated covariance matrix *C*, determined during burnin (computed sequentially 5 times), define the orthonormal eigenvectors *η_j_, Cη_j_* = *λ_j_η_j_*. Then propose a move in the two directions separately, *j* = 1, 2,

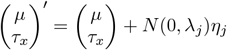

which has an acceptance probability,

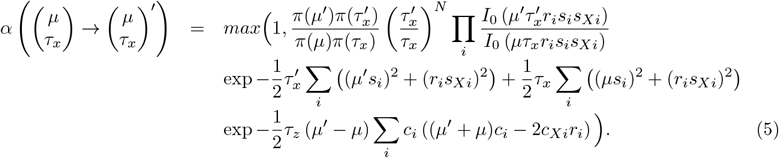

The priors enforce rejection of proposals that violate the positivity requirements, *μ* > 0, *τ_x_* > 0.

##### Precision 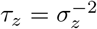

Using a conjugate Gamma prior Γ(*a_z_, k_z_*) we have the Gamma distributed update

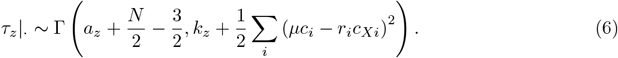

##### Hidden variable *θ_i_*

We use a random walk proposal. We used proposal 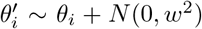, with step size *w* = 0.75 giving reasonable acceptance rates. The acceptance probability is,

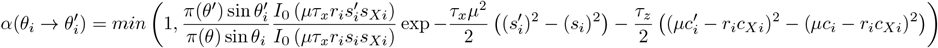

Since this is a symmetric random walk the proposal cancels.

##### Priors

The prior in *μ* is a truncated Gaussian (*μ* > 0) with large standard deviation, *μ* ~ *N*(60,100^2^). Priors for *τ_x_,τ_z_* are are weakly informative Gamma distributions, 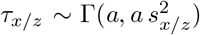 (shape and rate), with *a* = 10. This has mean precision *s*^−2^ and a relative standard deviation 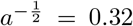. For *τ_x_,τ_z_* we used *s_x_* = 20, *s_z_* = 40 nm respectively when both flourophores are antibodies, and *s_x_* = 25, *s_z_* = 75 nm if one of them is GFP. Our posteriors are always tighter than these priors and typically have posterior means 20-30 nm for *σ_x_*, 40-55 nm for *σ_z_*.

We filter the data with the constraint |*X*| < 200 to limit the effect of the non-isotropy. Measured distances over 200 nm are clearly in error so such measurements are removed.

#### 1.5 MCMC algorithm performance on simulated data

We tested the algorithm on simulated data (see Figure below). Convergence was determined using the Gelman-Rubin diagnostic for multiple chains and determined converged if the corrected Gelman-Rubin statistic was below 1.1. We used 4 chains with chains initialized from the priors. The true parameters are accurately reconstructed, as seen in the following Figure.

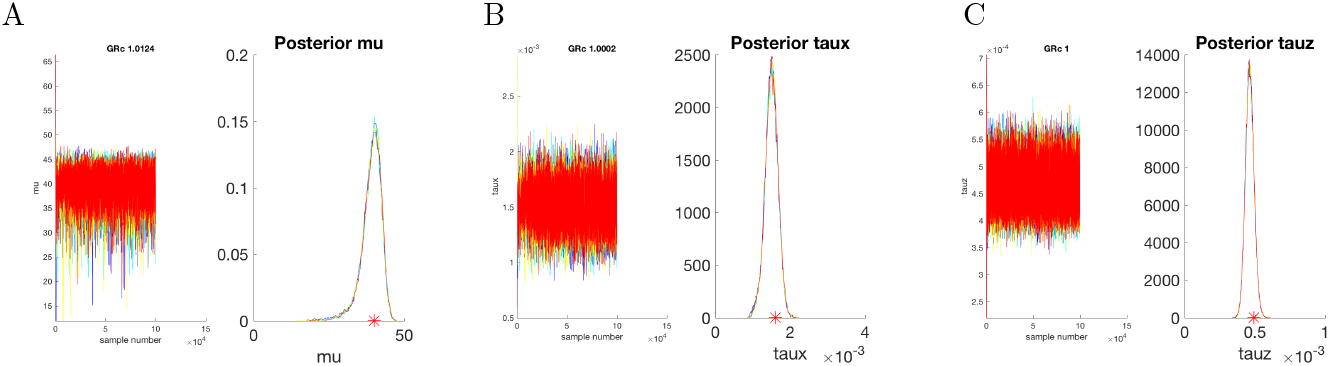
Performance on simulated data. Markov chains (4) and posterior distributions shown for distance *μ*, precision 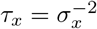, precision 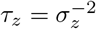. Chains shown in different colors. Corrected Gelman Rubin statistic (GRc) for that variable given at top of panel. True values shown on x-axis in red; *μ* = 40 nm, *σ_x_* = 25, *σ_z_* = 45 nm. MCMC runs of length 100000 including 50000 burnin. Priors for *τ_x, z_* had *a* = 3. Data set consists of 1000 simulated kinetochores.

#### 1.6 MCMC algorithm on experimental data

On all runs we used priors with *a* = 10. We used the Gelman-Rubin diagnostic for multiple chains to assess convergence, requiring that the corrected Gelman-Rubin statistic was below 1.1 on all three parameters *μ, τ_x_, τ_z_*. Burn-in was 50% of the run in all cases. Most data sets converged within 50000; for datasets that failed to converge within 50000 the run length was increased until convergenced attained. All histograms are based on a single run with 50000 samples (using subsampling at appropriate rates). The maximum run length (including burn-in) was 2000000 for the CenpC-to-Ndc80(C) distance in MC191 cell line under nocodazole treatment.

### 2 Kinetochore architecture simulations

#### 2.1 Structural data

We used kinetochore architectural information and structural data (crystalographic and EM), see Table below, to build a kinetochore simulation. The orientation of the Ndc80/Nuf2 calponin homogy (CH) domains with respect to the microtubule lattice, and the extending coiled coils were from Wilson-Kubalek et al. (2008). The Ndc80 hinge (also called loop or kink) was positioned 16 nm from the CH domains and allowed to bend and rotate given the intrinsic flexibility of the hinge and coiled coil that connects to the Spc24/Spc25 subunits (Maiolica et al., 2007; Huis in’t Veld et al., 2016; Scarborough et al., 2019; Wang et al., 2008). The Ndc80 hinge angle and the elevation angle of the NDC80 complex short arm (the coiled coil between Ndc80 hinge and Ndc80/Nuf2 CH domains) with respect to the microtubule lattice have been measured using purified complexes but not within the context of the intact kinetochore *in vivo*. So this information was not imposed.

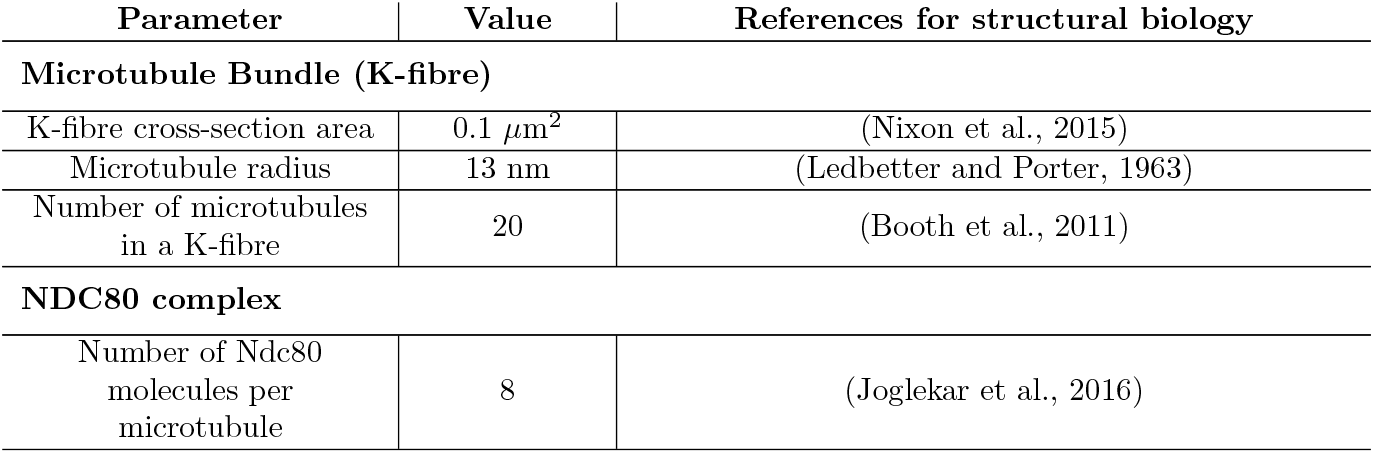
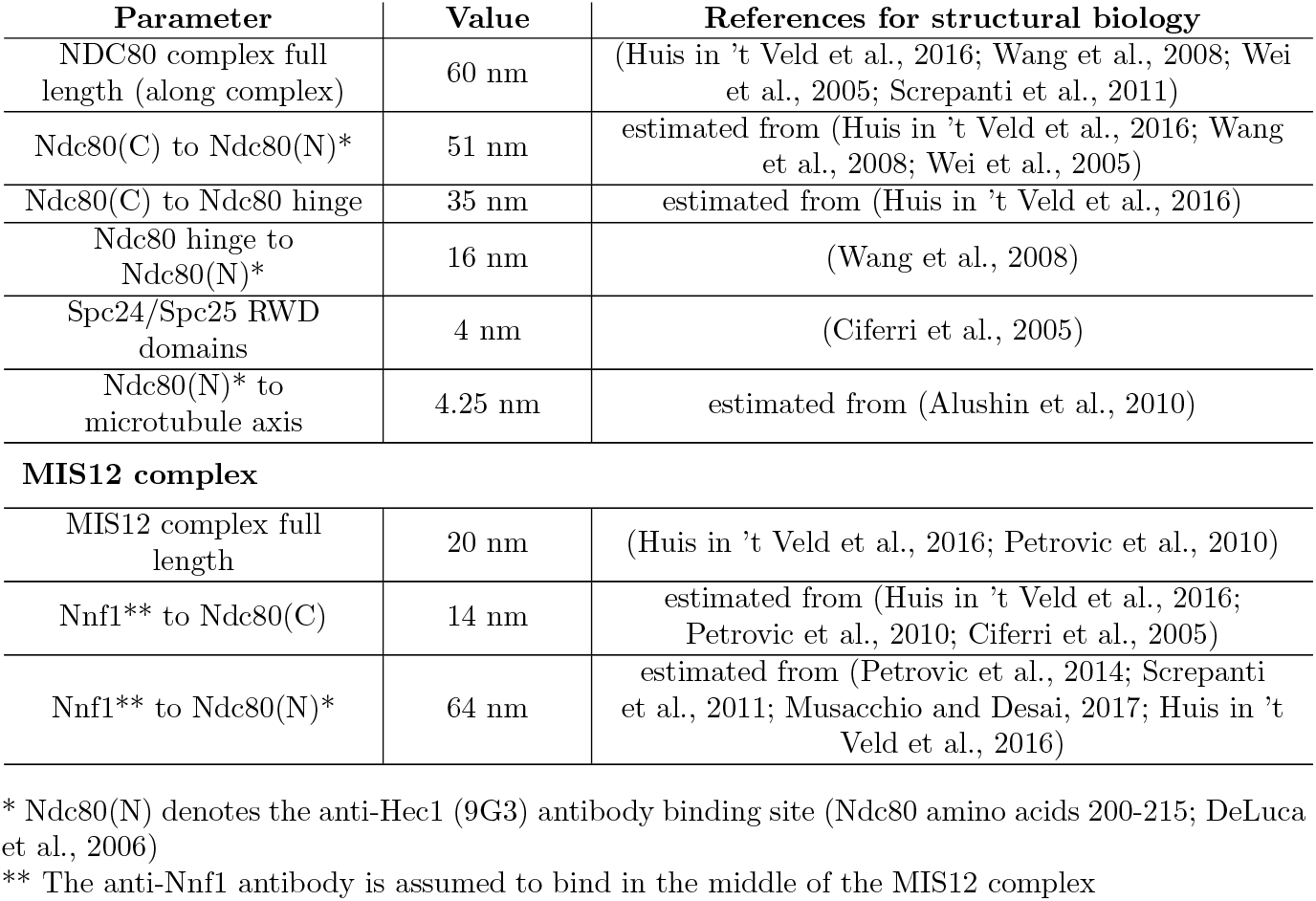

#### 2.2 Simulation

The simulation proceeds as follows. For parameters that are fixed from structural data, the values are described in Section 2.1. Parameters that are fitted to the Ndc80(N)-Ndc80(C)-CenpC triangle distances are indicated below (‘fitted’ in text, red color in schematic) and the fitting is described in Section 2.3. Note that variability in angles is set to 10° in absence of constraining data. See schematics below for graphical representation of the incorporated flexibility and annotations.

1. **Microtubule Bundle (K-fibre)**. The K-fibre is defined as a disc of diameter 360 nm, centred on the x-axis. Next, 20 microtubules (MT), radius 13 nm, are uniformly distributed within the K-fibre cross-section. Here, the microtubules lie parallel to the x-axis. If there are overlapping microtubules, all positions are rejected and the process is repeated.
2. **NDC80 complex.** The true mean of the Ndc80 attachment points is labelled x along the central axis of the K-fibre. Next, 8 NDC80 complexes are placed on each microtubule as follows: First, we determine a binding site along the MT (Gaussian, mean *x*, sd 25 nm) and rotate around the MT axis by angle *φ*_1_ (uniform in [0, 360°]). Next, the Ndc80(N) is positioned at 4.25 nm offset from the microtubule surface, considering the 9G3 antibody binding site. This results in 17.25 nm offset from the MT axis (13 nm MT radius). The NDC80 complex short arm is simulated by elevation at angle *φ*_1_ (Gaussian, fitted mean, sd 10°) relative to the microtubule axis, with the Ndc80 hinge oriented towards the inner kinetochore plate. The NDC80 complex short arm can tilt around the axis parallel to the x-axis going through Ndc80(N), angle *φ*_2_ (Gaussian, mean 0°, sd 10°), i.e. the NDC80 complex short arm and the microtubule axes would no longer lie in the same plane for non-zero tilt. The Ndc80 hinge is positioned at a fixed distance of 16 nm from Ndc80(N). The hinge angle between the NDC80 complex short arm and long arm (NDC80 complex coiled coil region between Spc24/Spc25 Head domains and Ndc80 hinge) is defined as *φ*_3_ (Gaussian, fitted mean, sd 10°). We also incorporate rotation of the long arm around the NDC80 complex short arm axis, angle *φ*_4_ (Gaussian, mean 0°, sd 10°). Ndc80(C) is positioned 35 nm from the Ndc80 hinge along the NDC80 complex long arm. Nnf1 is positioned a further 14 nm from Ndc80(C) along the NDC80 complex long arm axis towards the inner kinetochore. Finally, CenpC is positioned as described bellow.
3. **CenpC-Ndc80(N) axis**. Here, we use a model where the inner kinetochore (detected by the CenpC marker in this study) is off axis relative to the K-fibre. The inner-outer kinetochore axis is defined by elevating a line from the x-axis (pinned at *x*) at angle *φ*_5_ (Gaussian, fitted mean, sd) and rotating around the K-fibre axis by a random angle *φ*_6_ (uniformly distributed). The focus point *y* is positioned at a fixed distance (fitted) from *x* along this inner-outer kinetochore axis. CenpC is placed at a distance *u* (fitted) along the line joining Nnf1 to the focus point *y*; the focus will act to concentrate CenpC molecules relative to Nnf1. If *y* is at infinity there is no focusing, i.e. the spread of CenpC and Nnf1 will be identical.

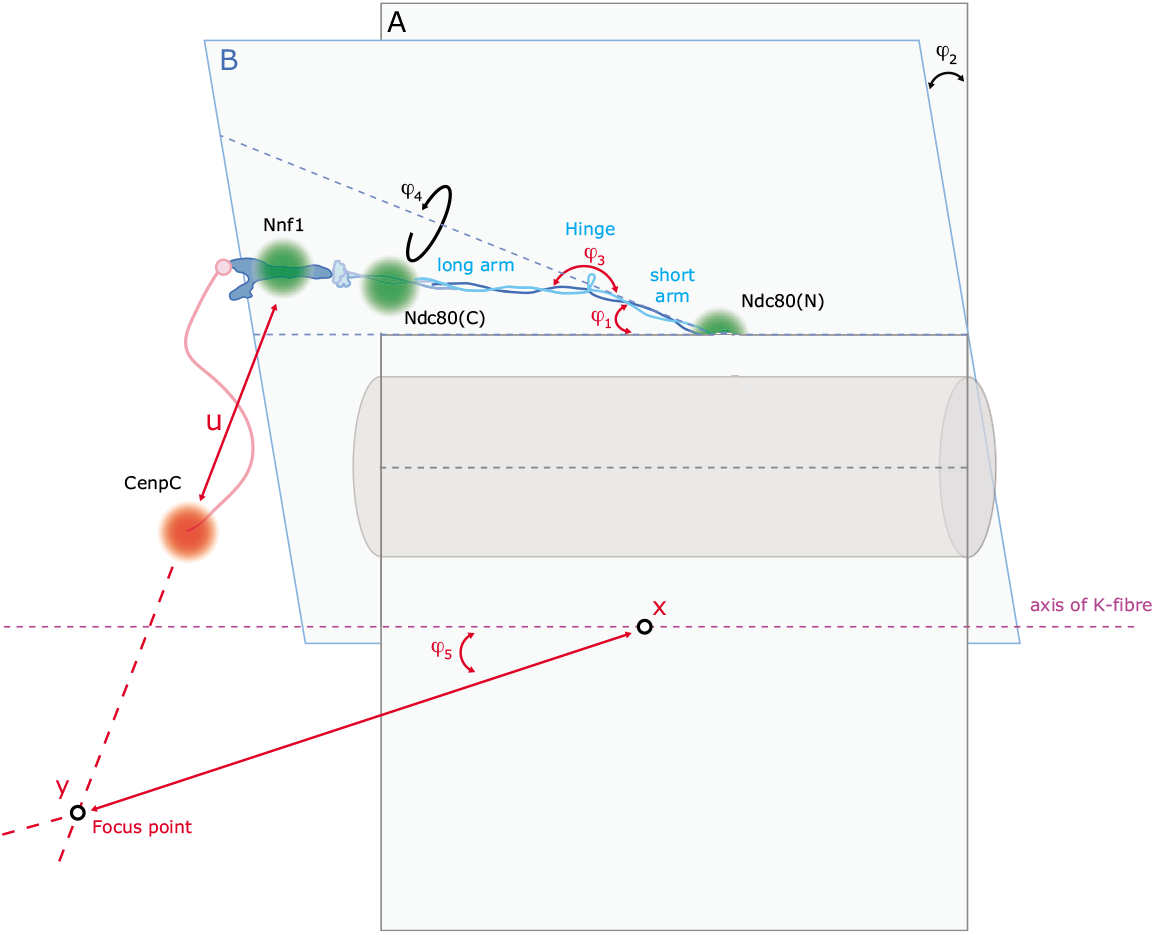
Geometrical setup for simulations of the CenpC-MIS12-NDC80 ensembles. Green and red dots represent the position of the indicated kinetochore proteins. A representative microtubule is shown in grey and its axis (black dotted line) lies on plane A. The purple dotted line indicates the K-fibre axis (x-axis). The NDC80 complex short arm lies on plane B and tilts (*φ*_2_) around the axis (light blue dotted line) that is parallel to the x-axis and goes through Ndc80(N). Fixed angle parameters used in the simulations are shown in black. Fitted parameters obtained from the simulations are instead displayed in red. For illustration only, the NDC80 MT attachment is shown vertically above the microtubule (*φ*_0_ = 0).

#### 2.3 Fitting of parameters. Optimisation

The measured △_*EC*_ distances in DMSO (cell line MC191) of 1. Ndc80(C)-to-Ndc80(N), 2. CenpC-to-Ndc80(N), and 3. CenpC-to-Ndc80(C), were used to fit six parameters: the focus point *y* (fitting distance from *x*, and mean, sd of angle *φ*_5_), the distance *u* from CenpC-to-Nnf1, the means of the short arm elevation angle and the Ndc80 hinge angle. We use a least-squares fitting method to minimise the difference between simulated distances in the triangle Ndc80(N)-Ndc80(C)-CenpC, and those observed. Specifically we minimise:

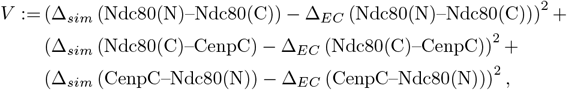

where the Δ_*sim*_ are the simulated distances, averaged over 20000 independently simulated kinetochores using the algorithm above.

The optimisation procedure is as follows: Starting from the current values of the 6 parameters a new set of these parameters is proposed based on a random walk (Gaussian) around the current values. The random walk has a drift term 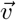 so that the algorithm continues to move in profitable directions, where the new drift vector is calculated for each jump and given by,

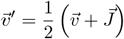

where 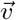 is the previous drift and 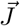 is the just accepted jump, if the proposal was accepted, and zero otherwise. The random walk has the following standard deviations: i) 50 nm for the |*x* – *y*| distance, ii) 2.5° for all angles except the mean of *φ*_5_, which was 5°. Larger step sizes were used early on to explore a larger part of the parameter space. A proposal is accepted if the cost function *V* is reduced. Otherwise, the proposal is rejected and a new proposal is attempted. As the simulations have many stochastic degrees of freedom, the Δ_sim_ are prone to fluctuations (despite the 20000 kinetochore sample size). To remove any bias, we re-evaluate *V* for a new, independent sample of 20000 kinetochores every time after an acceptance and every second time after a rejection. The process is completed if the results are considered close enough to the minimum of *V*, or if there is no improvement.

This method gives the following fit, reproducing the observed mean distances in the Ndc80(N)-Ndc80(C)-CenpC triangle to within 1 nm:

- angle *φ*_1_: mean 22.4°. This is within the range 20-60° given by Wilson-Kubalek et al. 2008.
- angle *φ*_3_: mean 203.5° (where 180° represents straight NDC80 complex conformation and angles >180° denote clockwise bending).
- distance |*x – y*|: 490 nm
- distance *u*: 37.9 nm
- angle *φ*_5_: mean 64.2°, sd 18.1°.

Here it needs to be noted that if additional constraints are imposed, a solution can still be found that fits the observed distances. For instance, imposing the elevation of the NDC80 complex short arm relative to the microtubule axis (*φ*_1_) to be 10°, sd 10° (restricted to be positive) gives a solution with Ndc80 hinge angle (*φ*_3_) mean of 177.3°. Therefore, there are nearby solutions that do not require the hinge angle to be above 180°.

### 3 Nematic order

Nematic order refers to the alignment of the molecules, alignment relative to the mean orientation axis. Orientation is determined relative to two flourophores. In the simulations nematic order *N* is defined as (| · | is Euclidean distance, < · > is average over sample),

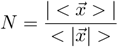

ie the length of the average vector to the average length of those vectors.

To compute the nematic order from experiment, we use the analogous formula,

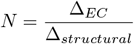

the ratio of the observed (mean) distance between two fluorophores and the expected structural distance Δ_*structural*_. Because each kinetochore is an ensemble, ͤ_*EC*_ is the (average) distance of the average vector over the ensemble.

